# Retinal Cx36 gap junctions in the inner and outer plexiform layers differentially control visual thresholds

**DOI:** 10.1101/2025.09.15.676372

**Authors:** Sam LaMagna, Eduardo Solessio

**Affiliations:** Center for Vision Research, Department of Ophthalmology and Visual Sciences, SUNY Upstate Medical University, Syracuse, New York 13210

**Keywords:** Gap junctions, visual thresholds, Cx36, temporal summation

## Abstract

Gap junctions are an important component of signal communication in sensory systems. In the retina, connexin 36 (Cx36) gap junctions integrate visual signals over space and time. However, we do not understand how Cx36 gap junctions in the inner- and outer-retina govern visual thresholds under dim light conditions. Here we address this question using transgenic mouse models, a state-of-the-art operant behavioral assay, and psychophysical modelling. Visual thresholds with Ganzfeld illumination decreased steeply (– 2 log/dec) with flash durations < 0.5 s, reaching an asymptotic value of *∼* 2 × 10^−4^photons/µm^2^/s with longer flashes. Temporal summation and photon noise modelled under Signal Detection Theory account for the observed thresholds. Removal of outer-retinal Cx36 lowers visual thresholds by 2.6-fold. Nested model analysis suggests that the probability of transmission of single photon signals between rods and bipolar cells increases in the absence of CX36 gap junctions between rods and cones. Removal of Cx36 in both the inner and outer retina increases visual thresholds by 16.5-fold, consistent with a loss of spatial-averaging and reduction of neural noise by Cx36 in the inner-retina. We conclude that both inner- and outer-retinal Cx36 control visual thresholds under dim light conditions, but in distinct ways. Outer-retinal Cx36 increases thresholds, perhaps in a necessary tradeoff that provides rod signals access to the alternative rod-cone pathway, whereas inner-retinal Cx36 decreases thresholds by increasing spatial-integration, more than compensating for outer retina losses.

## INTRODUCTION

Dark adapted, or ‘scotopic’, vision is highly sensitive, allowing us to detect singles-to-dozens of photons at a time (1, 2). Scotopic vision’s high sensitivity is a byproduct of its slow temporal kinetics (3, 4), high degree of spatial integration (5), and ability to keep low neural noise levels (6, 7). Connexin 36 (Cx36) gap junctions are considered essential for this remarkable sensitivity (8, 9), because they facilitate the spatiotemporal integration of neural signals in both the inner and outer synaptic layers of the retina (8, 10, 11).

In the inner synaptic layer, Cx36 is critical for function of the ON-branch of the rod ‘primary pathway’ (8, 9), where it electrically couples AII amacrine interneurons with ON cone bipolar cells (11–15). Further, inner retinal Cx36-mediated coupling between AII amacrine interneurons enhances the signal-to-noise ratio of dim light stimuli by averaging out spontaneous electrical activity (i.e., noise) and correlating upstream rod responses (i.e., signal) (16–18). This is important for signal fidelity in both the ON- and OFF-branches of the rod primary pathway (9). While these properties are known to underly the high sensitivity of the retina physiologically, how this translates to perceptual sensitivity is not yet known.

In the outer synaptic layer, Cx36-mediated coupling between rod and cone photoreceptors provides multiphoton rod signals with direct ‘piggy-back’ access to cone pathways, i.e. the rod ‘secondary pathway’ (8–10, 15, 19). However, the role of interphotoreceptor coupling with dim flashes eliciting single photon responses remains controversial. Cx36-mediated interphotoreceptor coupling reduces the amplitude of single photon responses in the rod cell (20–22). This has led some to predict that interphotoreceptor coupling is deleterious to visual sensitivity under single-photon conditions (21–23). However, spatiotemporal integration by Cx36-mediated coupling also reduces the spontaneous electrical activity that obscures single photon light responses (20, 24). This has led others to predict that the two effects cancel-out, and should leave visual sensitivity intact (20, 25).

To resolve these conflicting points of view, we took a ‘top-down’ approach, investigating the effects of inner and outer retinal Cx36 gap junctions on visual sensitivity at the level of behavior. We measured absolute visual thresholds (i.e., the lowest amount of light required to see) of two transgenic mouse lines: a Cx36 knockout (Cx36KO) mouse (8), which does not express Cx36 in either the inner or outer retina, and a photoreceptor specific-Cx36 knockout (Cx36XO) which does not express Cx36 between rods and cones, eliminating rod-cone and rod-rod coupling (10). Visual thresholds in response to full-field illumination were measured using a yes-no one-alternative forced choice (1AFC) behavioral assay wherein mice are trained to detect flashes of light (26). To further investigate the temporal integrative properties of Cx36 gap junctions, we obtained threshold measurements for a range of flash durations (0.06 s to 2 s) which were analyzed using a Theory of Signal Detection TSD (27)-based model of temporal summation developed here.

We found that both thresholds and psychometric function slopes were dependent on flash duration, and that our model could account for the flash duration dependence of thresholds. Cx36XO thresholds were 2.6-fold lower than their wild-type (WT) littermates, while Cx36KO thresholds were elevated 16.5-fold compared to their WT littermates. Further, using Gnat1 knockout (Gnat1KO) mice, which lack functional rods (28), we showed that Cx36KO mice do not use cone-mediated vision at absolute visual threshold. This suggests that visual thresholds in Cx36KO mice may be mediated by OFF rod-pathways. Our data and model of temporal summation suggest that inner retinal Cx36 lowers the absolute visual threshold by integrating rod signals, whereas outer retinal Cx36 may play a more subtle role, such as attenuating the single photon response (21, 23) or tuning signal transfer between rods and bipolar cells at threshold (7).

## RESULTS

### ABSOLUTE VISUAL THRESHOLDS DECREASE WITH FLASH DURATION

Visual thresholds in humans are inversely related to stimulus duration over short integration times, a phenomenon known as temporal summation (29–31). Here, we used an operant assay (26) to investigate the effect of flash duration on murine absolute visual thresholds for a range of flash durations (0.06 – 2 s) using a full-field flash stimulus (Fig.1A). To determine thresholds, we measured *d*^*′*^ psychometric functions over a 2.0 log unit intensity range at each flash duration. We defined the threshold intensity as the flash intensity corresponding to *d*^*′*^ = 1 in the psychometric functions. Figure 1B shows a family of psychometric functions for a single, representative, WT mouse (‘WT1’) taken at different flash durations. On the x-axis, intensity is expressed in terms of photons (ph) per µm^2^ per second (ph/µm^2^/s) at the retina. For each flash duration tested, *d*^*′*^ values rose monotonically with flash intensity, and were well fit by log-linear psychometric functions (Equation 3; see Methods) (26) (*R*^2^ > 0.9 at all flash durations). As flash duration increased, psychometric functions shifted leftwards, corresponding to a decrease in threshold intensity, up to 0.5 s. Threshold intensity appeared to plateau above 0.5 s as shown in the threshold vs duration (t.v.d.) plot (Fig.1C).

**Fig. 1.**
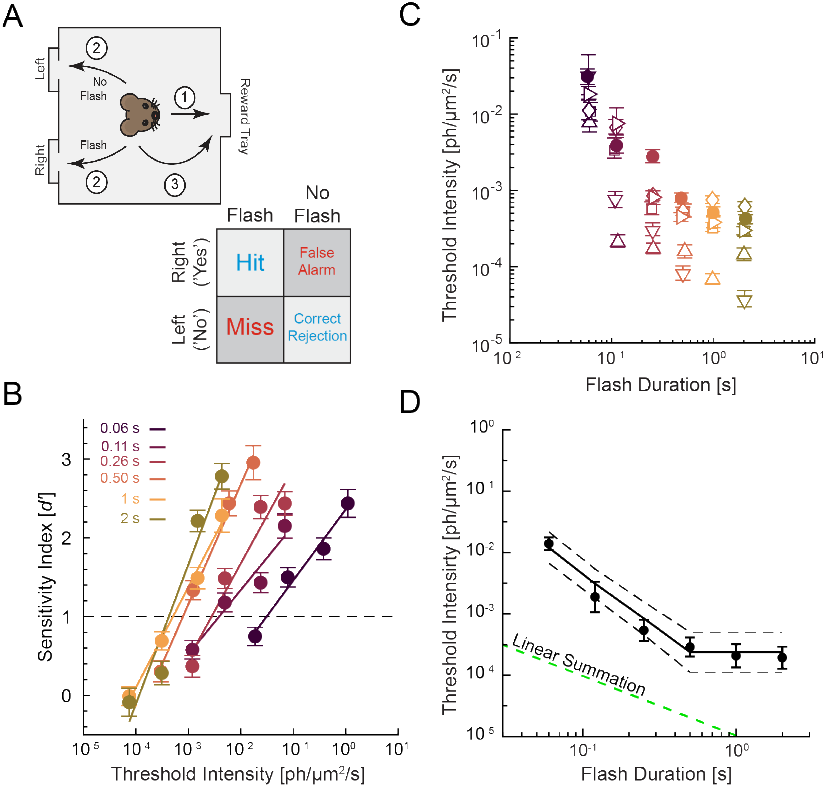
Temporal summation at absolute threshold. **A** Operant 1-AFC assay diagram and decision matrix. Mice initiate a trial by first visiting the reward tray (1); after an auditory count-down, the mouse visits either the left or right nose poke depending on whether a flash stimulus was presented (2); the mouse returns to the reward tray (3). If the mouse was correct (i.e., a “Hit” or “Correct Rejection”) it receives reward; if the mouse was incorrect (i.e., a “False Alarm” or “Miss”) it does not receive a reward. **B** *d*^*′*^ -psychometric functions for a single, representative mouse at different flash durations. This mouse is represented by a solid-colored symbol in panel C. Error bars are standard errors calculated via Equation 2 (see Methods). **C** Threshold versus duration (t.v.d.) plot of all WT mice (each represented by a different symbol). Error bars are 95% bias-corrected and accelerated (BCa) confidence intervals (CI) for thresholds obtained via non-parametric bootstrapping. **D** Averaged WT t.v.d. data with standard error, as well as the thresholds predicted by the best fit of our TSD model of temporal summation (black solid line) as well as 95% CI of model predictions (black dashed lines). Linear summation (green dashed line) is also shown to highlight the supra-linear nature of temporal summation observed and captured by our model.

The dependence of threshold on flash duration was found in all WT mice (Fig.1C-D and Fig.S1.1). We found that WT mice exhibited two regimes in their t.v.d. data. At shorter flash durations (below *∼* 0.5 s) thresholds decreased with flash duration, i.e., mice exhibited temporal summation (31). At longer flash durations thresholds ceased to rely on duration, indicating that summation was ‘complete’ (Fig.1C). This is more clearly seen in Figure 1D, which plots the average WT thresholds as a function of flash duration. This pattern was consistent with previous studies of temporal summation at absolute threshold in both human and amphibian subjects (29, 32–34). Interestingly, as duration increased, so did the steepness of the psychometric function slope (Fig.1B and Fig.S1.1). Mechanisms governing the slope of the *d*^*′*^ psychometric function are poorly understood, but are believed to involve factors such as uncertainty (35, 36), attention (37), and nonlinearities in stimulus transduction (38).

Threshold vs duration data are conventionally fit with Bloch’s law of linear temporal summation (30, 31, 39). However, this analysis provides no insight into the mechanisms governing how thresholds change with flash duration. We developed here an alternative model of temporal summation based on the Theory of Signal Detection (40) (i.e., the ‘Signal Detection Model’; see Fig.S1.2 and Methods for full derivation), which treats the retina as an array of independent detectors (e.g., ganglion cells), whose responses are integrated over time. The model fits our t.v.d. data better than linear temporal summation (see Fig.1D for comparison) and provides two free parameters needed for the discussion of our data: the effective number of rods for each detector, ‘*a*’, and an associated non-linearity with power coefficient ‘*b*’ which controls the steepness of the t.v.d. curves (see Methods). Fitting the model to WT data (Fig.1D) results in the following estimates: *a* = 3,119 ‘effective’ rods/detector [2,116; 4,591], and *b* = 0.30 [0.25; 0.34] (*R*^2^ = 0.41).

### GLOBAL DELETION OF CX36 RAISES THE ABSOLUTE VISUAL THRESHOLD

We first determined the impact of retinal Cx36 on the absolute visual threshold by measuring the absolute visual thresholds of Cx36KO (*n* = 8) and WT (n = 6) mice (Fig.2A). The Cx36KO mouse does not express Cx36 anywhere in the nervous system (8). Thus, it is feasible that changes to extra-retinal neural circuitry may be, in part, responsible for some changes in Cx36KO detection. Using a bright (*∼* 3.5 × 10^4^ph/µm^2^/s), long (2 s) flash stimulus, we ruled out this possibility; we found no differences in sensitivity indices (*d*^*′*^) between a subset of Cx36KO (*n* = 4) and WT (*n* = 3) mice (*t*_5_ = 0.55, *p* = 0.60). At these bright light levels, Cx36-dependent pathways (e.g., the ON branch of the rod primary pathway) were not used to drive visual behavior in either genotype, as was observed previously with electrophysiological recordings (19) and the processing of temporal contrast (41). Thus, for our 1AFC task, any observed differences in absolute visual threshold between Cx36KO and WT mice were due to the absence of retinal Cx36, specifically.

Like those of their WT littermates, *d*^*′*^ data of Cx36KO mice were well fit by log-linear psychometric functions at all durations (*R*^2^ > 0.9 for all Cx36KO mice; Figure S2.1). Cx36KO thresholds decreased with flash duration up to *∼* 0.5 s, after which thresholds remained constant (Fig.2B). However, averaged across durations, Cx36KO thresholds were elevated 16.5-fold relative to those of their WT littermates (Fig.2B and 2C). We used repeated measures (RM) linear mixed model (LMM) analysis to investigate the differences between WT and Cx36KO thresholds. RM LMM is interpreted similarly to a RM two-way ANOVA but allows for missing values and accounts for between-subject variability (see Methods). RM LMM analysis confirmed a 1.22 *±* 0.22 log_10_(ph/µm^2^/s) least square (LS) mean difference between WT and Cx36KO thresholds (*F*_5,59_ = 31.9, *p* < 1 × 10^−3^, *f* ^2^ = 0.19). RM LMM analysis also revealed a small-to-medium sized genotype *×* duration interaction effect (*F*_5,59_ = 2.6, *p* = 0.034, *f* ^2^ = 0.12; see Methods). This difference may be explained by a change in summation or a change in the non-linearity of the transduction channels (represented by model parameters *a* and *b*, respectively). Using nested model analysis, we found that only a is genotype-dependent (model probability = 83%; see Table 1 for model comparisons; see Fig.2C for best model predictions; and Fig.S2.3 for all model predictions): the WT-*a* was estimated to be 2,997 effective rods/detector [1,600; 5,526] and the KO-*a* was estimated to be 95 effective rods/detector [47; 167], a 31-fold difference. Per this model, genotype has no effect on *b*, which is estimated to be 0.26 [0.24; 0.29], similar to the value estimated previously for WT mice (Fig.1D). Thus, to some extent, the difference between thresholds of WT and Cx36KO mice can be explained by the effective number of rods per detector.

**Table 1.** Summary statistics of nested model fits for WT/Cx36KO genotype comparison. Note, the MSE of each model is included as a parameter for calculating the BIC.

| Model | Parameter Number | Independent Parameter | Shared Parameters | BIC | $\Delta\text{BIC}_i$ | $P(\text{Model})$ |
| --- | --- | --- | --- | --- | --- | --- |
| A (Full) | 5 | WT-*a*, WT-*b*, KO-*a*, KO-*b* | — | -98.21 | 3.24 | 0.17 |
| B | 4 | WT-*b*, KO-*b* | *a* | -49.77 | 51.68 | $5.01 \times 10^{-12}$ |
| C | 4 | WT-*a*, KO-*a* | *b* | -101.45 | 0 | 0.83 |
| D (Reduced) | 3 | — | *a*, *b* | -26.75 | 74.70 | $5.02 \times 10^{-17}$ |

### THRESHOLDS OF CX36KO MICE ARE NOT GOVERNED BY CONE MEDIATED SIGNALING

In the absence of Cx36, the thresholds of Cx36KO mice were either cone-mediated or rod-mediated via the OFF branch of the rod primary pathway (Fig.2A) (8, 9). To determine whether thresholds of Cx36KO mice were cone-mediated, we determined the absolute visual thresholds of Gnat1KO mice (*n* = 4) using our 1AFC task. Gnat1KO mice do not express the transducin *α* subunit, and thus lack functional rod vision, only cone-mediated vision (Fig.2A) (28). We hypothesized that if the thresholds of Gnat1KO mice were elevated relative to those of Cx36KO mice, we could conclude that thresholds of Cx36KO mice were not cone mediated. On the other hand, if the two genotypes exhibited similar thresholds, then we could not rule out cone mediated thresholds of Cx36KO mice.

Like *d*^*′*^ data of WT and Cx36KO mice, *d*^*′*^ data of Gnat1KO mice were well fit with log-linear psychometric functions at each flash duration tested (*R*^2^ > 0.9 for all Gnat1KO mice; Fig.S2.2). Also like the WT and Cx36KO genotypes, thresholds of Gnat1KO mice exhibited an initial inverse relationship with flash duration before plateauing (Fig.2B). Unlike the thresholds of WT and Cx36KO mice, thresholds of Gnat1KO mice decreased with flash intensity up until 1 s on average (Fig.2B). There was also more variability in threshold estimates between Gnat1KO mice compared to those of WT and Cx36KO mice (Fig.2B), which likely had to do with the smaller sample size of Gnat1KO mice. Notably, despite the larger variance and smaller sample size, analysis with RM LMM revealed a very large genotype dependent difference between the three genotypes (*F*_2,74_ = 220.1, *p* < 1 × 10^−4^, *f* ^2^ = 26.5). Specifically, there was a LS mean difference of 5.76 *±* 0.28 log_10_(ph/µm^2^/s) between WT and Gnat1KO thresholds across durations. This large difference is consistent with previous psychophysical investigations of the Gnat1KO mouse (42). Between thresholds of Cx36KO and Gnat1KO mice, there was a LS mean difference of 4.54 *±* 0.27 log_10_(ph/µm^2^/s). These findings suggest that the absolute thresholds of Cx36KO mice were not cone-mediated, leading us to infer that their absolute thresholds were mediated by signaling along the OFF branch of the rod primary pathway.

### PHOTORECEPTOR-SPECIFIC CX36 DELETION LOWERS ABSOLUTE THRESHOLDS

We next sought to determine the effects of photoreceptor Cx36 on the absolute visual threshold. To do this, we measured the absolute visual threshold of *HRGP*^*Cre*^::*Cx36*^*f/f*^ (Cx36XO; *n* = 7) mice and their *Cx36*^*f/f*^ littermates (WT^f/f^; *n* = 6) (10). Cx36XO cones do not express Cx36 (Fig.3A), which has been shown sufficient to eliminate rod-rod and rod-cone coupling (10). Like the other genotypes tested, log-linear psychometric functions fit both Cx36XO and WT^f/f^*d*^*′*^ data well across durations (*R*^2^ > 0.9, for all WT^f/f^and Cx36XO mice; Fig.S3.1 and S3.2). From Figure 3B, we can see that the thresholds of both WT^f/f^and Cx36XO mice decreased with flash duration up until *∼* 0.5 s, just as the previously discussed thresholds for WT and Cx36KO genotypes. Note, we collected data for most mice up to 1 s, as our WT, Cx36KO, and Gnat1KO data demonstrate that thresholds do not change from 1 s to 2 s. We tested two WT^f/f^and two Cx36XO at 2 s, to confirm that this pattern is also exhibited by these two genotypes. Thresholds of Cx36XO and WT^f/f^mice obtained for 2 s were not used in any further analysis.

**Fig. 2.**
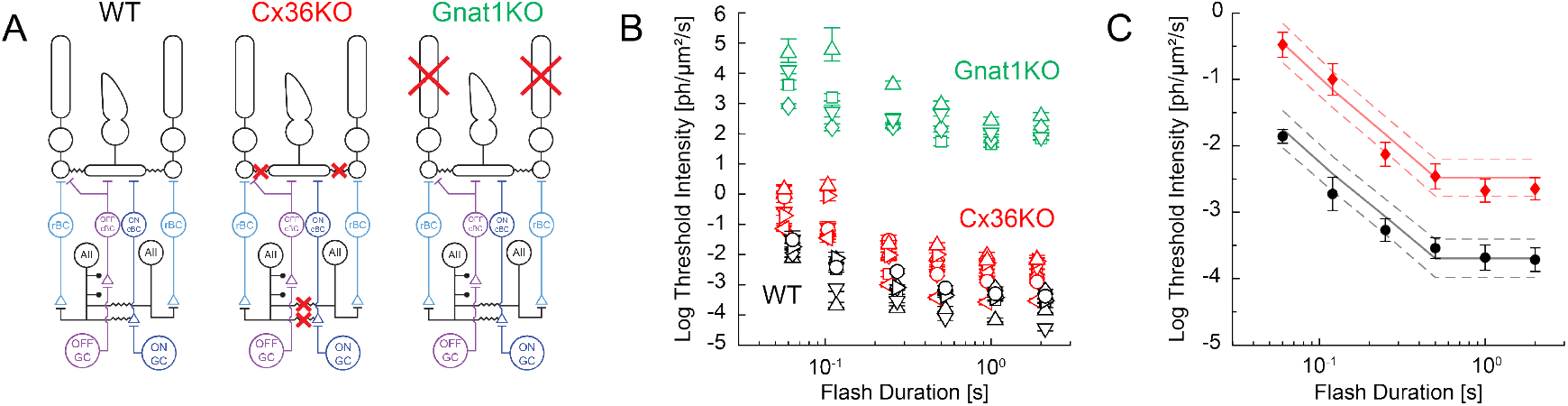
Absolute visual thresholds are elevated in the absence of inner and outer retinal Cx36 and rod phototransduction. **A** Wiring diagrams of WT, Cx36KO, and Gnat1KO retinae. In the WT mouse rod signals are transmitted via the ON branch of the rod primary pathway (rod *→* rod bipolar cell (rBC) *→* AII amacrine cell (AII) *→* ON cone bipolar cell (cBC) *→* ON ganglion cell (GC)). Cx36 gap junctions facilitate signal transfer from the ON cBC to the AII, as well as AII-AII and rod-cone coupling. In the Cx36KO mouse, Cx36 is removed from both the inner and outer retina, meaning that rod signals must traverse the OFF branch of the rod primary pathway (rod *→* rBC *→* AII *→* OFF cBC *→* OFF GC). Further, there is no AII-AII or rod-cone coupling. In the Gnat1KO mouse, there is no rod phototransduction, and thus all visual signals originate in the cones. **B** Log threshold intensity plotted as a function of flash duration for individual WT (black), Cx36KO (red), and Gnat1KO (green) mice. Error bars are 95% BCa confidence intervals for thresholds obtained via non-parametric bootstrapping. Note, data from WT and Cx36KO genotypes are slightly displaced horizontally for clarity. **C** Average log threshold intensity as a function of flash duration for WT (black circles) and Cx36KO (red diamonds) mice. Error bars are standard errors. Additionally, the model predictions for the best fit model (model C) are shown in solid lines; dashed lines are 95% CIs for the model predictions.

**Fig. 3.**
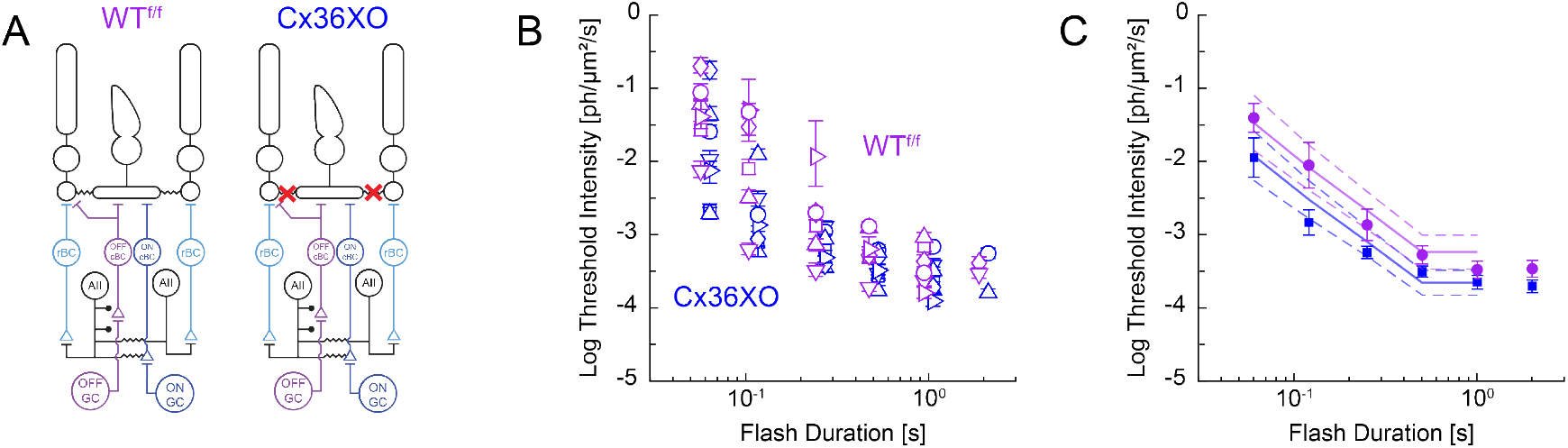
Absolute visual thresholds are lowered in the specific absence of outer retinal Cx36. **A** Wiring diagrams of WT^f/f^and Cx36XO retinae. In WT^f/f^retinae, Cx36 facilitates rod-cone coupling. In Cx36XO retinae, Cx36 is not expressed in cone cells, leaving rods and cones electrically decoupled. However, in both genotypes the rod primary pathway conveys visual signals at threshold. **B** Log threshold intensity plotted as a function of flash duration for individual WT^f/f^(blue) and Cx36XO (purple) mice. Error bars are 95% BCa CI for thresholds obtained via non-parametric bootstrapping. Note, data from both genotypes are slightly displaced horizontally for clarity. **C** Average log threshold intensity as a function of flash duration for WT^f/f^(blue circles) and Cx36XO (purple squares) mice. Error bars are standard errors. Additionally, the model predictions for the best fit model (model C) are shown in solid lines; dashed lines are 95% CIs for the model predictions.

There is more overlap in thresholds between genotypes than the previous experiments with WT and Cx36KO mice (Fig.3B). However, average Cx36XO thresholds were lower than average WT^f/f^thresholds at all durations, being more pronounced at shorter flash durations than at longer durations (Fig.3C). RM LMM analysis revealed that there is a 0.42 *±* 0.18 log_10_(ph/µm^2^/s) LS mean difference between thresholds of Cx36XO and WT^f/f^mice (*F*_1,44_ = 5.55, *p* = 0.02, *f* ^2^ = 0.07). Nested model analysis (Methods and Table 2) suggests that the different thresholds may arise from a 4.8-fold difference in the ‘effective’ spatial summation parameter (model probability = 70%; best model predictions in Fig.3C; all model predictions in Fig.S3.3), where WT^f/f^-*a* was estimated to be 1,280 effective rods/detector [537; 2,469] and the XO-*a* was estimated to be 4,336 effective rods/detector [2,081; 7,806], with a shared *b* = 0.29 [0.26; 0.33]. These results show that there are important qualitative differences in the role that retinal Cx36 plays in scotopic vision of Cx36KO and Cx36XO mice.

**Table 2.** Summary statistics of nested model fits for WT^f/f^/Cx36XO genotype comparison. Note, the MSE of each model is included as a parameter for calculating the BIC.

| Model | Parameter Number | Independent Parameter | Shared Parameters | BIC | $\Delta\text{BIC}_i$ | $P(\text{Model})$ |
| --- | --- | --- | --- | --- | --- | --- |
| A (Full) | 5 | $\text{WT}^{\text{f/f}}\text{-a}, \text{WT}^{\text{f/f}}\text{-b}, \text{XO-a}, \text{XO-b}$ | — | -85.32 | 1.77 | 0.29 |
| B | 4 | $\text{WT}^{\text{f/f}}\text{-b}, \text{XO-b}$ | $a$ | -76.20 | 10.89 | $3.01 \times 10^{-3}$ |
| C | 4 | $\text{WT}^{\text{f/f}}\text{-a}, \text{XO-a}$ | $b$ | -87.09 | 0 | 0.70 |
| D (Reduced) | 3 | — | $a, b$ | -78.48 | 8.61 | $9.44 \times 10^{-3}$ |

## DISCUSSION

In this study, we combined a novel operant behavioral assay with Signal Detection modeling to investigate how Cx36 gap junctions govern absolute visual thresholds. We have shown that both inner- and outer-retinal Cx36 govern the absolute visual threshold, but in different ways. Selective loss of interphotoreceptor Cx36-mediated coupling decreases absolute visual thresholds, whereas loss of Cx36 in both the outer and inner retina increases visual thresholds relative to controls. This result has important implications to our understanding of the mechanisms that set absolute visual thresholds.

### RECEPTOR CX36 GAP JUNCTIONS INCREASE ABSOLUTE VISUAL THRESHOLDS

Our first major result is that the thresholds of Cx36XO mice were 2.6-fold lower than their WT^f/f^littermates. Photoreceptor gap junctions in mice are mediated by Cx36 (43) and selective knockout in cones eliminates cone-cone, cone-rod, as well as rod-rod coupling (10). Rod-cone coupling provides rod signals with a portal to the cone pathways (21, 23, 44, 45) extending the rod-driven responses well into the upper mesopic range (19). Rod-rod coupling, whether direct (24, 45–49) or indirect by way of cones (rod-cone-rod) (21, 50, 51), has been proposed to play a critical role in the transmission of single photon information from rods to bipolar cells in dim lights.

Quantitative analysis suggests that rod-rod coupling can be beneficial (improve signal to noise ratio in response to uniform illumination) if the number of activated rods is larger than the ‘effective’ number of coupled rods, where ‘effective’ refers to the number of rods within the immediate coupled network. However, coupling can be detrimental (reduction in the signal to noise ratio) with dimmer, sparse illumination, such that a single rod is activated within its ‘effective’ coupling radius (21, 22). More recent analysis poses that rod responses in mouse are independent of the degree of coupling (25). We find that in response to short flash durations (0.06 s), the absolute visual thresholds in mouse occur at 0.01 ph/µm^2^/s, or equivalently 8 × 10^−4^ R*/rod, which activates less than one out of every 1,000 rods. In this case, the probability of activating more than one rod within the effective coupling area is negligible and in line with the suboptimal summation conditions outlined above. Therefore, the reduction in absolute visual thresholds that we observe in Cx36XO relative to those in control mice is in line with the physiological models that predict interphotoreceptor coupling to be deleterious to the absolute psychophysical threshold (21–23).

Our detection model provides insights into an alternative mechanism underlying the reduction in threshold in Cx36XO mice relative to their control siblings. As shown in Eq. 5.2, the parameter *a* reflects the combined effects of spatial summation, in terms of the number of rods in the receptive field of the detector (*a*^*′*^), and the efficiency of photon response transmission (*γ*). In the mammalian retina, a rod bipolar cell will receive input from *∼* 20 rods (5, 52, 53), each generating noise that, when summed linearly, will obscure true rod single photon responses (54, 55). To mitigate this problem, a synaptic thresholding mechanism is tuned to preferentially transmit large single photon responses while attenuating small responses and cellular noise (7). Given that the murine rod-to-rod bipolar synapse is estimated to transmit *∼*25% of single photon responses (7), we can conservatively assign *γ* a value of 0.25 in equation 5.2. Thus, *a* = 1,287 effective rods/ detector for WT^f/f^control mice, can be understood in terms of an *a*^*′*^*≈* 5,000 rods/detector when 25% thresholding is considered. Interestingly, 5,000 rods/detector approximates the convergence values estimated for ON *α* ganglion cells (16), and is in line with the receptive field area of the ON *α* ganglion cells (56, 57) when assuming a rod density of 437,000 rods/mm^2^ (58).

Since Cx36 deletion is limited to the photoreceptors in the Cx36XO mouse, it is unlikely that this change in *a* reflects a change in spatial summation at the level of the ganglion cells, but could represent a change in the efficiency of single photon response transmission, *γ*. There are two possible ways photoreceptor Cx36 modulates *γ*: by reducing the signal-to-noise ratio of single photon responses (21, 23) or by fine-tuning the thresholding mechanism indirectly. Rod-cone coupling may alter the local membrane potentials at the terminals, given that rods and cones have different resting potentials (-34 vs -45 mV, respectively (20)). This would lead to a change in the gain of the calcium and voltage-dependent release of glutamate, thus repositioning the thresholding nonlinearity (7, 55, 59) and the efficiency of single photon response transmission.

Our findings raise an important question: what benefit is gained from interphotoreceptor coupling that is worth the reduced sensitivity? As indicated above, rod-cone coupling provides rod signals with a portal to the cone pathways (21, 23, 44, 45) extending the rod-driven responses well into the upper mesopic range (19). Physiological models demonstrate that the signal-to-noise ratio can be enhanced by coupling when multiple rods in a coupled network are illuminated (20, 25, 48, 60), which would enhance object detection in the dark. It has also been suggested that photoreceptor coupling helps to extend the operating range of the rod cell, which is predicted to enhance contrast detection in brighter lights (23, 48, 61).

### INNER RETINAL CX36 DECREASE ABSOLUTE VISUAL THRESHOLD

We found that there is, on average, an approximately 16.5-fold difference in thresholds between WT and Cx36KO mice. Our model attributes the difference to a change in the *a* parameter. Unlike the Cx36XO mouse, Cx36 is not expressed anywhere in the Cx36KO retina (8). This makes it more difficult to attribute the change in *a* to any retinal mechanism or circuit. However, given our finding that eliminating Cx36 expression in the outer retina lowers thresholds, we can infer that loss of Cx36 in the inner retina is specifically responsible for the higher Cx36KO thresholds.

In the inner plexiform layer, Cx36 is predominantly expressed by AII amacrine cells, where it mediates AII-AII, AII-ON cone bipolar cell, and AII-*α* ganglion cell coupling (8, 11–13, 62). Specifically, it is AII-expressed Cx36 that is responsible for the high sensitivity of ON ganglion cells (9). Thus, it is possible that the decreased a value in the Cx36KO mouse is due to loss of AII-expressed Cx36, which enhances the signal-to-noise ratio of rod signals by averaging out baseline noise (16, 17) and mediating the pooling of multiple, independent, rod signals (16, 17, 63).

### OFF ROD-SIGNALING AT ABSOLUTE VISUAL THRESHOLD IN CX36KO

Cx36KO mice can only see via the OFF-rod pathways or ON/OFF-cone pathways (8, 9, 19). Electrophysiological recordings show that the cone-mediated thresholds of Cx36KO ON ganglion cells are elevated *∼* 1,000-fold relative to WT (8, 19). In contrast with these findings, we only observed a 16.5-fold elevation of psychophysical thresholds of Cx36KO mice relative to WT mice. Our behavioral observations are more consistent with the 10-fold difference in sensitivity between WT ON ganglion cells and Cx36KO OFF ganglion cells (9). Further, we found that the absolute visual threshold of Cx36KO mice is several orders of magnitude lower than that of cone-mediated vision. Prior evidence shows that mice can use the OFF pathway for detecting suprathreshold cone-mediated light increments (64). However, it is known that at scotopic threshold WT mice use the ON pathway to encode light increments (6, 57, 65). Our results are consistent with the notion that in the absence of the ON pathway, mice can use OFF pathway encoding of increment stimuli at their absolute threshold.

### TEMPORAL SUMMATION AT THE MURINE ABSOLUTE VISUAL THRESHOLD

While a critical duration of 0.2 s has been widely assumed in murine studies of scotopic vision (e.g., (16, 42, 66) this value is not experimentally established for the mouse. Here, we show experimentally that mice integrate rod signals over a period of *∼* 0.5 s. This is 5-times the estimated critical duration for human scotopic vision (0.1 s) (29). Furthermore, our t.v.d. data does not follow the typical linear summation observed in human studies (29, 32). Indeed, prolonging flash duration from 0.06 s to 0.11 s, results in a 0.87 log unit difference in the thresholds of WT mice; the difference predicted by linear summation is 0.26 log units. Similarly, the difference in threshold from 0.11 s to 0.26 s is 0.54 log units, which is larger than the 0.36 log unit prediction of linear summation.

Deviations from linear summation are not uncommon at absolute threshold, where sublinear, or ‘partial’, summation is known to occur (29, 33, 67). However, supralinear summation, as seen here, has only been observed in humans under certain conditions; Dawson & Harrison observed supra-linear temporal summation when using a large, photopic stimulus against a dark background (68), similar to our use of a Ganzfeld stimulus. Thus, supralinear temporal summation may be a consequence of large visual stimuli which engage multiple detection mechanisms working in parallel as predicted by our double-integrator model. This deviation from linearity is a surprising result because the number of channels activated at visual threshold is expected to increase linearly with the duration of the flash (Eq.11). We found that under these conditions, the supra-linear temporal summation is governed by the parameter b, the power coefficient of the transduction stage of each channel. The best fitting value for *b* was equal to *∼* 0.3, which results in a compressive nonlinearity that turns out to be a good approximation to the logarithmic growth of the *d*^*′*^ -psychometric functions in WT, Cx36KO and Cx36XO mice.

## CONCLUSION

In summary, our findings clarify the impact of Cx36 on the absolute visual threshold. Our results demonstrate that Cx36-mediated electrical coupling of rods and cones in the outer retina is deleterious to sensitivity, most likely as a tradeoff for signal transmission at higher light levels. Meanwhile, Cx36 in the inner retina is critical for sensitivity. Nested model analysis attributes this to modulation of spatial integration in the inner retina and efficiency of transmission of single photon information at the rod to bipolar cell synapse in the outer retina

## METHODS

### EXPERIMENTAL MODEL AND SUBJECT DETAILS

#### Mouse strains and husbandry

We used the following strains of mice (3-9 months in age) maintained on a C57BL/6J background: *Cx36*^ࢤ/-^(Cx36KO) (8), *Gnat1*^ࢤ/-^(Gnat1KO) (28), *HRGP*^*Cre*^::*Cx36*^*f/f*^(Cx36XO) (10). The Cx36KO mouse is a global Cx36 knockout, where only Cx36-independent retinal pathways are functional, including OFF-rod pathways and cone pathways (8, 9, 19). The Cx36XO mouse is a cone-specific Cx36 knockout where Cx36-mediated electrical coupling is only disrupted between rods and cones, leaving the primary pathway intact (10, 19). The Gnat1KO mouse lacks the transducin *α*-subunit required for phototransduction in rods, leaving cones as the only functional photoreceptor within the stimulus range of our experiments (28).

We used eight Cx36KO mice, and five wildtype (*Cx36*^+/+^; WT) littermates obtained from heterozygous (*Cx36*^+/-^) breeder pairs. An additional C57BL/6J WT mouse from a separate litter was added for a total of six WT mice used as controls. We applied a similar strategy to generate Cx36XO mice. We obtained seven Cx36XO mice, and six *Cx36*^*f/f*^(WT^f/f^) littermates that were used as controls, by crossing *Cx36*^*f/f*^mice with *HRGP*^*Cre*^::*Cx36*^*f/f*^mice. The *HRGP*^*Cre*^ mouse expresses Cre specifically in cones (69), and crossing it with the *Cx36*^*f/f*^mouse removes Cx36 specifically between rods and cones (10). By ensuring that experimental and control animals came from the same litter, we minimized variability due to litter-specific differences in behavior. Four Gnat1KO mice were obtained by setting homozygous *Gnat1*^ࢤ/-^breeder pairs. Because the differences between Gnat1KO and WT thresholds were so large, we used WT mice instead of litter-matched control mice.

Genotypes for all mice were confirmed via PCR. Mice were maintained on a 14-hour light/ 10-hour dark cycle and dark-adapted overnight for at least 8 hours before each experiment. All tests were performed during subjective daytime hours, ZT = 4h to 8h. Our mice were bred on a C57BL/6J background, wherein melatonin synthesis is impaired (70). Thus, we did not expect circadian regulation of rod-cone coupling (50). Overnight dark adaptation ensures conductive rod-cone gap junctions in WT and WT^f/f^mice.

To maintain motivation for learning and performing the operant behavioral task, each mouse was kept on a food-restricted schedule (resulting in body weight between 80-90 percent of expected weight) as described by Umino et al., 2018 (71) and LaMagna et al., 2024 (26). We also determined whether the animals were lethargic or showed any signs of dehydration. If the animals’ body weight dropped to <80% of their normal body weight, we supplemented their diet with additional dry food on a daily basis. We progressively increased the daily amount of dry food that the underweight mouse received, in 0.25 g increments, until the mouse reached the target >80% expected weight. All procedures in this study were approved by the Institutional Animal Care and Use Committee at SUNY Upstate Medical University and were conducted in accordance with National Academy of Sciences’ *Guide for the Care and Use of Laboratory Animals* and in compliance with the Association for Research in Vision and Ophthalmology *Statement for the Use of Animals in Ophthalmic and Vision Research*.

### METHODS DETAILS

#### Operant behavioral assay to test absolute visual thresholds in mice

To determine absolute visual thresholds in mice we applied an operant conditioning assay (26) at various flash durations. Tests were performed using a control and conditioning system (Lafayette Instruments) consisting of eight experimental chambers controlled by ABET II software (Lafayette Instruments). The user-defined program scheduled the tests, recorded the responses, and presented the reinforcements. Each chamber consisted of a reward tray and two equidistant nose-pokes located on the opposite side. Light stimulation was presented by a programmable LED (Luxeon) placed overhead. The LEDs had central emission at 496 nm and a half-maximal bandwidth of 28 nm as measured with a spectral radiometer (Photo Research, SpectraColorimeter, model PR-650). Flash intensities were controlled by pulse width modulation and neutral density filters. Light diffusers were used to keep luminance levels between all surfaces of the chamber (ceiling, walls, and floor) uniform (0.3 log-unit maximal difference) as measured with a Graseby S370 optometer equipped with a photometric filter and 15°Lumilens.

Here we describe the assay briefly (for details see (26). The experimenter was blinded to genotypes during the conditioning, training, and testing of mice. Mice were trained to perform a yes-no, one-alternative forced choice (1AFC) task to discriminate the presence or absence of a flash in each trial (Fig. 1A). During the task, mice self-initiated trials by visiting a reward tray. Upon initiation, an auditory ‘countdown’ cue began, consisting of three 0.1 s high-frequency tones. 0.1 s after the last countdown tone a solid tone was presented concurrent with either a flash stimulus or no-flash stimulus. Following the presentation of the stimulus, mice were trained to visit one of two equidistant nose-pokes located opposite to the reward tray, dependent on the stimulus presented. Following the nosepoke visit mice returned to the reward tray; if the response was correct, they were rewarded with a small amount of commercially available Ensure Nutrition Drink (5 to 10 µL, Abbott Laboratories). The solid tone terminated when the mouse returned to the reward tray, or, in the absence of a response, after 14 s when the trial was aborted. Each completed trial had four possible outcomes (Fig.1A): (1) correctly visiting the right nose-poke on flash trials, i.e. a Hit; (2) correctly visiting the left nose-poke on no-flash trials, i.e. a Correct Rejection; (3) incorrectly visiting the right nose-poke on a no-flash trial, i.e. a False Alarm (FA); (4) incorrectly visiting the left nose-poke on a flash trial, i.e. a Miss.

We used the 1AFC task to measure absolute visual threshold so that we could utilize bias-independent measures of visual detection. Response bias, or an observer’s *a priori* predisposition to respond in one way or another, is known to distort common measures of visual detection, such as percent correct; in turn, this can lead to misleading estimates of threshold intensities (27, 72, 73). The 1AFC task allows us to measure both the probability of Hits (i.e., the Hit rate; *p*_*H*_) and the probability of FAs (i.e., the FA rate; *p*_*F*_), which we can use to calculate a bias-independent measure of detection, the sensitivity index (*d*^*′*^):

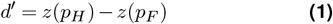

Where *z*(*·*) is the inverse of the standard normal distribution (72). Using *d*^*′*^ gives bias-independent measures of thresholds in mice (26, 71). For some mice, under certain conditions, there were instances of perfect detection (i.e., *p*_*H*_ = 1 and *p*_*F*_ = 0), and instances where the stimulus was completely undetectable (i.e., *p*_*H*_ = 0), which makes the calculation of *d*^*′*^ impossible. To get around this, we used a correction following Hautus, (74) where 0.5 is added to all *p*_*H*_ and *p*_*F*_. This method does not introduce statistical bias into *d*^*′*^ estimates (72, 74), and will converge to the ‘true’ population *d*^*′*^ within *∼* 400 trials (74) (as used here). Because our *d*^*′*^ values are based on pooled trials, we calculated the *d*^*′*^ standard error (SE) as (72):

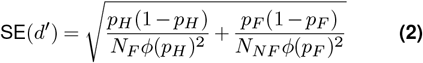

Where *ϕ*(*·*) is the height of the normal density function at *z*(*·*), and *N*_*F*_ and *N*_*NF*_ are the numbers of flash, and no flash, trials respectively.

#### Conditioning & training protocols

Mice were trained to complete the 1AFC task via a two-stage conditioning and training protocol (26). Briefly, mice first completed a five-phase operant conditioning protocol using a bright flash, 2 s in duration. In phase 1 mice are trained to associate the reward tray with reward. In phases 2 and 3, mice are trained to associate visiting the left nose-poke with no-flash stimuli, and the right nose-poke with flash stimuli, respectively. In phase 4, mice are presented with consecutive blocks of three flash and three no-flash trials. During this phase, if a mouse responds incorrectly the trial is repeated until the mouse responds correctly; these are termed corrective trials. Phase 4 continues until the mouse responds correctly 70% of the time. In phase 5, flash and no-flash trials are presented at random, with corrective trials still being in use. Phase 5 continues until the mouse responds correctly 80% of the time for three consecutive days. Genotype had no effect on the time required to complete the conditioning protocol (*F*_4,27_ = 1.74, *p* = 0.17, oneway ANOVA).

After completing operant conditioning, mice underwent a three-phase training protocol to move mice from long-duration stimuli to short-duration stimuli. Specifically, mice stabilized their day-to-day *d*^*′*^ values (defined as a coefficient-of-variation (CV) *≤* 20% over four days) for a flash stimulus with a duration (Δ*t*) of 2 s. This was then repeated for flashes with Δ*t* = 0.25 s, and 0.06 s, consecutively. All flashes were 400 R*/rod/s in intensity. Note, one Gnat1KO mouse (Gnat1KO-2 ‘*△*’ in Figure 2B) was especially desensitized compared to its littermates, and struggled to complete single-intensity training at Δ*t* = 0.06 s. To make sure Gnat1KO-2 had enough time to complete the entire sequence, we started measuring thresholds of Gnat1KO at Δ*t* = 2 s without completing single-intensity training at Δ*t* = 0.06 s. With this one exception, all mice completed the same single-intensity training protocols. Genotype had no effect on the time required to complete single-intensity training (*F*_4,25_ = 2.19, *p* = 0.1, one-way ANOVA excluding Gnat1KO-2).

### Measuring absolute visual thresholds

Each multi-intensity experimental session consisted of 400 trials of four randomly presented intensity levels over a 1.7-2.0 log-unit range, with no-flash trials randomly interspersed. The experimental session followed a single-intensity warm-up session, where the flash intensity was chosen to be in the middle of the range presented during the experimental session. These sessions consisted of at least 100 corrective trials and lasted 0.5-1 hr.

After mice demonstrated a stable *d*^*′*^ for Δ*t* = 0.06 s, mice began completing multi-intensity experiments at relatively bright flash stimuli (*∼* 1-115 ph/µm^2^/s). Intensity was then gradually lowered, usually by 0.6 log-units, until performance stabilized near absolute threshold. We defined stability as CVs for individual *d*^*′*^ values *≤* 45% over three to four days, which is the upper limit for stabilized psychometric functions. ‘Near absolute threshold’ was defined as the intensity range where *d*^*′*^ values systematically crossed 1. Data that met our stability near threshold criteria were pooled across sessions and fit with the following *d*^*′*^ psychometric function:

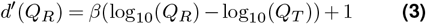

Where *Q*_*R*_ is the flash intensity in terms of photon flux density at the retina (see below), *β* is the psychometric function slope, and *Q*_*T*_ is the threshold intensity, i.e. the intensity where *d*^*′*^ = 1. Psychometric functions were fit via non-linear least squares regression as described below.

After obtaining thresholds with the 0.06 s stimulus, we then determined their thresholds for an additional 5 durations: 0.11 s, 0.26 s, 0.5 s, 1 s, and 2 s. We used the same stability near threshold criteria as for the 0.06 s stimulus for all other durations. To average out any learning effects that may occur as mice progress through flash durations, we randomly assigned mice to complete multi-intensity experiments for the additional flash durations in either an ascending (0.11 s 2 *→* s) or descending (2 s*→* 0.11 s) sequence. Mice were tested between 3-9 months of age. One WT mouse (WT-5, ‘▽’ in Fig.1C and 2B) was not capable of completing the entire sequence before it aged out. For the purpose of model fitting, thresholds at 2 s were more useful than those at 1 s, thus we skipped Δ*t* = 1 s for WT-5, and obtained a threshold at Δ*t* = 2 s instead.

#### Retinal illumination

Luminance incident from the chamber walls and ceiling was measured with a Graseby S370 optometer equipped with a 15°Lumilens. We found that the luminance of the surfaces was within approximately 0.3 log units of one another, consistent with an isoluminant visual environment. To determine retinal irradiance, we first made radiometric readings using the Graseby S370 in radiometric mode and by positioning the photodiode in the location of the mouse cornea directed at the back of the chamber, the nose-poke-side of the chamber, and ceiling of the chamber. These values were averaged together to arrive at the typical irradiance within the chamber.

Radiometric measurements were converted to corneal irradiance (*Q*_*C*_(*λ*)) at a given wavelength *λ* = 497 nm (central emission wavelength of the LED stimulus), in terms of photons per µm^2^ per second (ph/µm^2^/s) at the cornea. *Q*_*C*_(*λ*) was converted to retinal irradiance (*Q*_*R*_(*λ*)), in terms of ph/µm^2^/sat the retina, via Lyubarsky et al., 2004 (75):

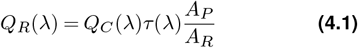

Where *τ* (*λ*) is the fraction of photons transmitted through the ocular media, *A*_*P*_ is the area of the dark-adapted pupil, and *A*_*R*_ is the area of the retina. *τ* (*λ*) was set to 0.7, *A*_*R*_ to 18 mm^2^ (75) and *A*_*P*_ to 4 mm^2^ (76). We estimated the average number of photoisomerizations, *Q*_*p*_(*λ*), (in R*/rod/s), by multiplying *Q*_*R*_(*λ*) by the end-on collecting area of the rod cell (*a*_*c*_(*λ*)), which is 0.87 µm^2^ (75):

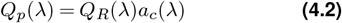

### QUANTIFICATION AND STATISTICAL ANALYSIS

#### A Signal Detection Theoretic model of temporal summation

Our model consists of two integration stages. In the first stage, multiple (*N*) parallel signal detectors (40) respond independently to the flash stimulation (Fig.S1.2). The effective number of photons ‘seen’ by a single detector over an integration time (Δti) is given by:

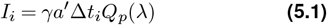

Where *Q*_*p*_(*λ*) is the average photoisomerization rate per rod (Eq.4.2), *a*^*′*^ is the total number of rods within the receptive field area of a single detector, and *γ* is a scaling factor which accounts for the proportion of photoisomerization responses that are successfully transmitted to the detector. For convenience we introduce the parameter, *a* defined as:

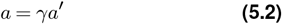

Such that a represents the ‘effective’ number of rods in a receptive field after taking *γ* into account.

The detector is characterized by a nonlinear transducer with a power coefficient *b*, such that

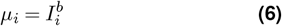

We further assumed that the sensitivity of the detector is limited by external (*σ*_*E*_) and internal ‘dark’ (*σ*_*D*_) noise sources (Fig.S1.3A). The external noise is known by the nature of the stimulus. Under dim light levels photon counting is a Poisson process (1, 77), where the standard deviation is defined as the square root of the mean intensity. The external noise level after the nonlinear transformation is:

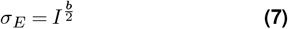

Internal noise reflects baseline neural activity, which is comprised of both peripheral photoreceptor noise (78) and central neural noise (79). We make the simplifying assumption that central noise is negligible relative to photoreceptor noise (80). Assuming a rod collecting area of 0.87 µm^2^, we adopt an internal noise value of *D* = 0.01 R*/rod/s (42) over an integration window Δ*t*_*i*_, such that the input to the channel detectors is given by:

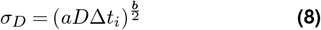

For independent noise sources, the total variance at the channel detector is given by:

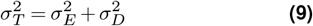

To determine *d*^*′*^ for a single detector 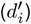, we systematically varied *µ* and the corresponding *σ*_*T*_ over the dynamic range of the stimulus (10^−5^ to 10^1^ R*/rod/s) to generate *signal + noise* and *noise* distributions. As a first approximation we assumed that these functions were described by Gaussian distributions, where the noise distribution had a mean *m* = 0 and *σ*_*D*_ given by equation 8, while the signal + noise distributions had a mean *µ*_*i*_ given by equation 6 and *σ*_*T*_ given by equation 9. We then determined the *p*_*F*_ and *p*_*H*_ by setting a variable criterion and estimating the areas under the noise and signal + noise distributions above the criterion, respectively. Using this approach, the area under the noise distribution provides a measure of *p*_*F*_ while the area under the signal + noise distribution provides a measure of *p*_*H*_. Repeating this procedure for multiple criteria produced plots of *p*_*H*_ vs *p*_*F*_, also known as receiver operating characteristic (ROC) curves. We plotted the ROC curves on *z*-coordinates and found that as flash stimuli become brighter, extrinsic noise predominates (Fig.S1.3B) and the z-score ROCs derived for the model exhibit slope values (*s*) < 1 (Fig.S1.3B) that increase with stimulus intensity, as is expected for Poisson distributed noise (81–83). Under these conditions, the *d*^*′*^ may not eliminate decision bias (72). Instead, the slope of the *z*-score ROC is used to estimate 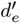, an alternate measure of sensitivity that is independent of decision bias (72):

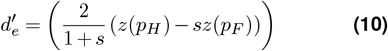

However, in general, we found that 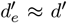 when *d*^*′*^ = 1 (Fig.S1.3B and S1.4), confirming that *d*^*′*^ is an accurate measure of sensitivity at threshold.

The second stage of the model uses additive summation to derive the total *d*^*′*^ (27, 40, 84), where the number of detectors increases linearly with flash duration. It is straightforward to see that as flash duration, Δ*t*, increases, the number of detectors responding to the stimulus (*N*), each with an integration time Δ*t*_*i*_, increases linearly according to 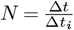. Thus, we can use additive summation to derive the total *d*^*′*^ corresponding to *N* as (27, 40, 84):

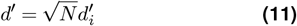

We apply Equation 11 to the *d*^*′*^ -psychometric function for a single detector, generating a family of *d*^*′*^ -psychometric functions for different values of *N*. Figure S1.2C shows a family of *d*^*′*^ -psychometric functions estimated this way for *N* = 1, 2, 4, 8, 8, and 8 (see justification for these values below). The resulting functions reproduce features of the *d*^*′*^ -psychometric functions observed in WT mouse (Fig.1). As the number of detectors increased, the psychometric functions shifted to the left although they did not show major changes in slope (Fig.S1.3C). In analogy to the mouse behavioral data, we defined thresholds as the intensities where the modelled *d*^*′*^ -psychometric functions intersect the line *d*^*′*^ = 1, and the slopes as the tangents at the threshold points (represented with thick lines in Fig.S1.3C). FigureS1.3D shows model predictions as a function of systematical changes in the parameters *a, b*, and the dark noise (*D*). Both *a* and *D* uniformly scale thresholds across duration and thus set the location of the t.v.d. curve along the threshold intensity axis. Parameter *b* controls the slope of the t.v.d. functions, implicating it directly in temporal integration. A linear model with the same parameter values does not produce the steep t.v.d. plots predicted by the TSD model (Fig.S1.3D-iv).

The last step required to set up the model was to specify the integration times of each stage. We tentatively assigned the integration time, Δ*t*_*i*_, of the detectors in the first stage a value Δ*t*_*i*_ = 0.06 s in analogy to the time constant of ON *α* retinal ganglion cells relaying visual information at threshold levels (85). The integration time of the second stage can be inferred from the data. The duration at which thresholds begin to plateau, i.e., the critical duration, is approximately 0.5 s (Fig.1). Using the critical duration as a measure of the integration limit of the second summation mechanism (40, 86), we limit the integration time of the second stage to 0.5 s, which, for Δ*t*_*i*_ = 0.06 s, results in a maximum *N* = 8.

#### Nested model analysis

To gain further insight into observed differences between mutant and wild-type mice, we conducted nested model analysis. For each wildtype mutant pair (i.e., WT-Cx36KO and WT^f/f^-Cx36XO) data were fit with four different nested models, each consisting of a different combination of parameters that were either genotype-dependent (i.e., estimated separately) or genotype-independent (i.e., estimated jointly) (see Fig.S2.3, Fig.S3.3, and Tables 1 and 2). The parameters in question included the summation factor, *a*, and the transducer nonlinearity, *b*. The most constrained nested model (model ‘D’) did not consider any genotype-dependent parameter and thus consisted of only the *a* and *b* parameters. The least constrained model (model ‘A’), considered both parameters as genotype-dependent, and thus consisted of four free parameters: an *a* for both genotypes and a *b* for both genotypes. The remaining nested models included a model where only *b* was genotype-dependent (model ‘B’), with separate *b* parameters for each genotype and shared *a* (three free parameters) and a model where only *a* was genotype dependent (model ‘C’), with an *a* for each genotype and a shared *b* (three free parameters).

#### Information theoretic model selection

The informationtheoretic approach to model selection overcomes limitations observed in other commonly used model selection approaches (87, 88). In model selection, we seek to choose the model with the least number of parameters (i.e., minimizing bias in parameter estimates) while also minimizing the mean square error (i.e., minimizing variance in parameter estimates) (87). Information theoretic model selection techniques are well-suited for this problem. Bayesian information criteria (BIC) is a measure of model goodness-of-fit that penalize extra parameters (87), which BIC is better suited for explanation than the AIC (89, 90). Thus, we based our model selection on BIC values. The BIC was calculated from least-squares fits via:

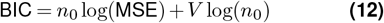

Where *n*_0_ is the number of observations, MSE is the mean square error of the fit, and *V* is the number of free model parameters; a lower BIC value corresponds to a better model fit. Note, for all model fits, the MSE is included as a free parameter (87).

The BIC also allows us to utilize the posterior model probability (*P* (model)), which provides an intuitive quantification of model fit (87, 91). The *P* (model) is the probability that a given model is correct, given the set of models and that the prior model probabilities of the set are all equal (87, 91). The posterior model probability is calculated as (91):

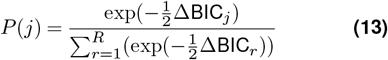

Where ‘*j*’ is the model index, ΔBIC_*j*_ is the difference between the BIC of model ‘*j*’ and the lowest BIC in the set, and *R* is the total number of candidate models.

#### Statistical analysis

All statistical analysis was carried out using SAS software (SAS Institute) and MATLAB software (Math Works). For all statistical tests, *α* = 0.05. To obtain psychometric function parameters, Equation 3 was fit to each mouse’s individual *d*^*′*^ data, for each Δ*t*, via nonlinear least squares regression using the `NLIN` procedure in SAS.

The model was fit using `lsqcurvefit` (MATLAB) to determine the coefficients a and b that best fit the threshold data. Upper and lower bounds of *a* and *b* were set at 0 – 20,000 and 0 – 1, respectively. Initial values were *a* = 3,000 and *b* = 0.5. Minimal step tolerance was set to 1 × 10^−6^. Nested models included data from both genotypes being compared (i.e., WT & Cx36KO or WT^f/f^ & Cx36XO) and the number of parameters was systematically reduced as described above. Bootstrapping via resampling with replacement was used to generate distributions of model parameters, from which 95% confidence intervals were determined (92) of parameter estimates. Nonparametric bootstrapping was used to calculate bias corrected and accelerated (BCa) confidence intervals of threshold estimates (26, 84) using custom SAS code. Individual experiments were simulated by randomly sampling a binomial distribution specified by the *p*_*H*_ or *p*_*F*_ and the number of flash or no-flash trials, respectively. Simulated *p*_*H*_ and *p*_*F*_ were then used to calculate simulated *d*^*′*^ values, from which we obtained simulated thresholds.

To test for differences in thresholds between genotypes we used repeated-measures (RM) linear mixed model (LMM) analysis using the `MIXED` procedure (93). The RM LMM is similar to the RM ANOVA, except it can take intra-subject variability into account, increasing statistical power. We modelled log threshold intensity with genotype, flash duration, and genotype *×* flash duration interactions as fixed effects, and a random intercept term for each mouse with an unstructured covariance matrix. Model fits were obtained using restricted maximum likelihood (REML). The inclusion of the random intercept term with unstructured covariance provided a significantly better fit than a fixed-effects only model for all comparisons (*p ≤* 0.001, null model likelihood ratio test), justifying the LMM approach. Effect sizes for fixed effects were reported as Cohen’s *f* ^2^ calculated according to Selya et al., 2012 (94). According to Cohen’s guidelines: 0.02 *≤ f* ^2^*≤* 0.15 is a small-to-medium effect, 0.15 *≤ f* ^2^*≤* 0.35 is a medium-to-large effect, and *f* ^2^ *≥* 0.35 is a large effect (95). One-way ANOVAs and t-tests were carried out using the `ANOVA` and `TTEST` procedures, respectively (93).

## REFERENCES

1. Selig Hecht, Simon Shlaer, and Maurice Henri Pirenne. Energy, Quanta, and Vison. The Journal of General Physiology, 1941(1926): 819–840, 1942.

2. B Sakitt. Counting every quantum. The Journal of Physiology, 223(1): 131–150, 1972. ISSN 14697793. doi: 10.1113/jphysiol.1972.sp009838.

3. J. D. Conner. The temporal properties of rod vision. The Journal of Physiology, 332(1):139–155, nov 1982. ISSN 14697793. doi: 10.1113/jphysiol.1982.sp014406.

4. Andrew Stockman, Lindsay T Sharpe, Eberhart Zrenner, and Knut Nordby. Slow and fast pathways in the human rod visual system: electrophysiology and psychophysics. Journal of the Optical Society of America A, 8(10): 1657, 1991. ISSN 1084-7529. doi: 10.1364/josaa.8.001657.

5. Peter Sterling, Michael A. Freed, and Robert G. Smith. Architecture of rod and cone circuits to the on-beta ganglion cell. Journal of Neuroscience, 8(2): 623–642, 1988. ISSN 02706474. doi: 10.1523/jneurosci.08-02-00623.1988.

6. Petri Ala-Laurila and Fred Rieke. Coincidence detection of single-photon responses in the inner retina at the sensitivity limit of vision. Current Biology, 24(24): 2888–2898, 2014. ISSN 09609822. doi: 10.1016/j.cub.2014.10.028.

7. Greg D. Field and Fred Rieke. Nonlinear signal transfer from mouse rods to bipolar cells and implications for visual sensitivity. Neuron, 34(5): 773–785, 2002. ISSN 08966273. doi: 10.1016/S0896-6273(02)00700-6.

8. Michael R. Deans, Béla Völgyi, Daniel A. Goodenough, Stewart A. Bloomfield, and David L. Paul. Connexin36 is essential for transmission of rod-mediated visual signals in the mammalian retina. Neuron, 36(4):703–712, nov 2002. ISSN 08966273. doi: 10.1016/S0896-6273(02)01046-2.

9. Béla Völgyi, Michael R. Deans, David L. Paul, and Stewart A. Bloomfield. Convergence and Segregation of the Multiple Rod Pathways in Mammalian Retina. Journal of Neuroscience, 24(49):11182–11192, feb 2004. ISSN 0270-6474. doi: 10.1523/JNEUROSCI.3096-04.2004.

10. Nange Jin, Zhijing Zhang, Joyce Keung, Sean B Youn, Munenori Ishibashi, Lian-Ming Tian, David W Marshak, Eduardo Solessio, Yumiko Umino, Iris Fahrenfort, Takae Kiyama, Chai-An Mao, Yanan You, Haichao Wei, Jiaqian Wu, Friso Postma, David L Paul, Stephen C. Massey, and Christophe P. Ribelayga. Molecular and functional architecture of the mouse photoreceptor network. Sci. Adv, 6 (28):7232–7240, jul 2020. ISSN 2375-2548. doi: 10.1126/SCIADV.ABA7232.

11. Stephen L Mills, Jennifer J. O’Brien, Wei Li, John O’Brien, and Stephen C Massey. Rod pathways in the mammalian retina use Connexin 36. Journal of Comparative Neurology, 436(3): 336–350, 2001. ISSN 00219967. doi: 10.1002/cne.1071.

12. Karin Dedek, Konrad Schultz, Mario Pieper, Petra Dirks, Stephan Maxeiner, Klaus Willecke, Reto Weiler, and Ulrike Janssen-Bienhold. Localization of heterotypic gap junctions composed of connexin45 and connexin36 in the rod pathway of the mouse retina. European Journal of Neuroscience, 24(6): 1675–1686, 2006. ISSN 0953816X. doi: 10.1111/j.1460-9568.2006.05052.x.

13. Andreas Feigenspan, Barbara Teubner, Klaus Willecke, and Reto Weiler. Expression of neuronal connexin36 in AII amacrine cells of the mammalian retina. Journal of Neuroscience, 21(1): 230–239, 2001. ISSN 02706474. doi: 10.1523/jneurosci.21-01-00230.2001.

14. E. Brady Trexler, Wei Li, Stephen L. Mills, and Stephen C. Massey. Coupling from AII amacrine cells to ON cone bipolar cells is bidirectional. Journal of Comparative Neurology, 437(4):408–422, sep 2001. ISSN 00219967. doi: 10.1002/cne.1292.

15. E. Brady Trexler, Wei Li, and Stephen C Massey. Simultaneous contribution of two rod pathways to AII amacrine and cone bipolar cell light responses. Journal of Neurophysiology, 93(3): 1476–1485, 2005. ISSN 00223077. doi: 10.1152/jn.00597.2004.

16. Felice A. Dunn, Thuy Doan, Alapakkam P. Sampath, and Fred Rieke. Controlling the gain of rod-mediated signals in the mammalian retina. Journal of Neuroscience, 26 (15):3959–3970, 2006. ISSN 02706474. doi: 10.1523/JNEUROSCI.5148-05.2006.

17. Robert G. Smith and Noga Vardi. Simulation of the AII amacrine cell of mammalian retina: Functional consequences of electrical coupling and regenerative membrane properties. Visual Neuroscience, 12(5): 851–860, 1995. ISSN 14698714. doi: 10.1017/S095252380000941X.

18. Noga Vardi and Robert G. Smith. The AII amacrine network: Coupling can increase correlated activity. Vision Research, 36(23): 3743–3757, 1996. ISSN 00426989. doi: 10.1016/0042-6989(96)00098-3.

19. Nange Jin, Lian Ming Tian, Iris Fahrenfort, Zhijing Zhang, Friso Postma, David L. Paul, Stephen C. Massey, and Christophe P. Ribelayga. Genetic elimination of rod/cone coupling reveals the contribution of the secondary rod pathway to the retinal output. Science Advances, 8(13): 1–12, 2022. ISSN 23752548. doi: 10.1126/sciadv.abm4491.

20. Nange Jin, Alice Z. Chuang, Philippe J. Masson, and Christophe P. Ribelayga. Rod electrical coupling is controlled by a circadian clock and dopamine in mouse retina. Journal of Physiology, 593(7):1597–1631, apr 2015. ISSN 14697793. doi: 10.1113/jphysiol.2014.284919.

21. Robert G. Smith, Michael A. Freed, and Peter Sterling. Microcircuitry Architecture of the Dark-Adapted Cat Retina: Functional Architecture of the Rod-Cone Network. The Journal of Neuroscience, 6(December):3505–3517, 1986.

22. M Tessier-Lavigne and D Attwell. The effect of photoreceptor coupling and synapse nonlinearity on signal : noise ratio in early visual processing. Proceedings of the Royal Society of London, 234: 171–197, 1988.

23. Eric P Hornstein, Jan Verweij, Peter H Li, and Julie L Schnapf. Gap-junctional coupling and absolute sensitivity of photoreceptors in macaque retina. Journal of Neuroscience, 25(48): 11201–11209, 2005. ISSN 02706474. doi: 10.1523/JNEUROSCI.3416-05.2005.

24. T. D. Lamb and E. J. Simon. The relation between intercellular coupling and electrical noise in turtle photoreceptors. The Journal of Physiology, 263(2): 257–286, 1976. ISSN 14697793. doi: 10.1113/jphysiol.1976.sp011631.

25. Nange Jin, Munenori Ishibashi, Zhijing Zhang, Stephen C. Massey, and Christophe P. Ribelayga. The impact of rod/cone electrical coupling on the rod single-photon response - a computational study. Investigative Opthalmology & Visual Science, 64(8), 2023.

26. Sam LaMagna, Yumiko Umino, and Eduardo Solessio. Signal Detection Theoretic Estimates of the Murine Absolute Visual Threshold Are Independent of Decision Bias. eNeuro, 11(10): 1–17, 2024. doi: 10.1523/ENEURO.0222-24.2024.

27. David M. Green and John A. Swets. Signal Detection Theory and Psychophysics. John Wiley & Sons, Ltd, New York, 1966.

28. Peter D. Calvert, N. V. Krasnoperova, A. L. Lyubarsky, T. Isayama, M. Nicoló, B. Kosaras, G. Wong, K. S. Gannon, R. F. Margolskee, R. L. Sidman, Edward N. Pugh, Clint L. Makino, and J. Lem. Phototransduction in transgenic mice after targeted deletion of the rod transducin α-subunit. Proceedings of the National Academy of Sciences of the United States of America, 97(25): 13913–13918, 2000. ISSN 00278424. doi: 10.1073/pnas.250478897.

29. Horace B. Barlow. Temporal and spatial summation in human vision at different background intensities. The Journal of Physiology, 141(2): 337–350, 1958. ISSN 14697793. doi: 10.1113/jphysiol.1958.sp005978.

30. A.M. Bloch. Experiences sur la vision. Comptes Rendus de la Societe de Biologie, 37(28): 493–495, 1885.

31. Andrew B Watson. Temporal Sensitivity, 1986.

32. L.T. Sharpe, C. Fach, and K. Nordby. Temporal Summation in the Achromat. Vision Research, 28: 1263–1269, 1988. ISSN 21644527. doi: 10.1080/02673843.1989.9747652.

33. P. Zuidema, H. Verschuure, Maarten A. Bouman, and J. J. Koenderink. Spatial and Temporal Summation in the Human Dark-Adapted Retina. Journal of the Optical Society of America, 71(12): 1472–1480, 1981. ISSN 00303941. doi: 10.1364/JOSA.71.001472.

34. Charlotte Haldin, Soile Nymark, Ann Christine Aho, Ari Koskelainen, and Kristian Donner. Rod phototransduction determines the trade-off of temporal integration and speed of vision in dark-adapted toads. Journal of Neuroscience, 29(18): 5716–5725, 2009. ISSN 02706474. doi: 10.1523/JNEUROSCI.3888-08.2009.

35. Denis G. Pelli. Uncertainty explains many aspects of visual contrast detection and discrimination. Journal of the Optical Society of America A, 2(9): 1508, 1985. ISSN 1084-7529. doi: 10.1364/josaa.2.001508.

36. Christopher W. Tyler. Why We Need to Pay Attention to Psychometric Function Slope. In Vision Scienceand its Applications, pages 240–243, Santa Fe, New Mexico United States, 1997.

37. Leonid L Kontsevich and Christopher W Tyler. Distraction of attention and the slope of the psychometric function. Journal of the Optical Society of America A, 16(2): 217–222, 1999.

38. Frederick A.A. Kingdom, Alex S. Baldwin, and Gunnar Schmidtmann. Modeling probability and additive summation for detection across multiple mechanisms under the assumptions of signal detection theory. Journal of Vision, 15 (5):1–16, 2015. ISSN 15347362. doi: 10.1167/15.5.1.

39. Kristian Donner. Temporal vision: Measures, mechanisms and meaning. Journal of Experimental Biology, 224(15), aug 2021. ISSN 14779145. doi: 10.1242/jeb.222679.

40. Andrei Gorea and Christopher W. Tyler. New look at Bloch’s law for contrast. Journal of the Optical Society of America A, 3(1): 52, 1986. ISSN 1084-7529. doi: 10.1364/josaa.3.000052.

41. Rose Pasquale, Yumiko Umino, and Eduardo Solessio. Rod photoreceptors signal fast changes in daylight levels using a CX36-independent retinal pathway in mouse. Journal of Neuroscience, 40(4): 796–810, 2020. ISSN 15292401. doi: 10.1523/JNEUROSCI.0455-19.2019.

42. Frank Naarendorp, Tricia M Esdaille, Serenity M Banden, John Andrews-Labenski, Owen P Gross, and Edward N. Pugh. Dark light, rod saturation, and the absolute and incremental sensitivity of mouse cone vision. Journal of Neuroscience, 30(37):12495–12507, sep 2010. ISSN 02706474. doi: 10.1523/JNEUROSCI.2186-10.2010.

43. Munenori Ishibashi, Joyce Keung, Catherine W Morgans, Sue A. Aicher, James R. Carroll, Joshua H. Singer, Li Jia, Wei Li, Iris Fahrenfort, Christophe P. Ribelayga, and Stephen C. Massey. Analysis of Rod/Cone Gap Junctions from the Reconstruction of Mouse Photoreceptor Terminals. eLife, 11, 2022. ISSN 2050084X. doi: 10.7554/eLife.73039.

44. Norianne T. Ingram, Alapakkam P. Sampath, and Gordon L. Fain. Voltage-clamp recordings of light responses from wild-type and mutant mouse cone photoreceptors. Journal of General Physiology, 151(10), 2019.

45. David M. Schneeweis and Julie L Schnapf. Photovoltage of rods and cones in the macaque retina. Science, 268(5213): 1053–1056, 1995. ISSN 00368075. doi: 10.1126/science.7754386.

46. D. Attwell, M. Wilson, and S. M. Wu. A quantitative analysis of interactions between photoreceptors in the salamander (Ambystoma) retina. The Journal of Physiology, 352(1): 703–737, jul 1984. ISSN 14697793. doi: 10.1113/jphysiol.1984.sp015318.

47. Peter B. Detwiler, Alan L. Hodgkin, and P. A. McNaughton. Temporal and spatial characteristics of the voltage response of rods in the retina of the snapping turtle. The Journal of Physiology, 300(1):213–250, mar 1980. ISSN 14697793. doi: 10.1113/jphysiol.1980.sp013159.

48. Peter H Li, Jan Verweij, James H Long, and Julie L Schnapf. Gap-junctional coupling of mammalian rod photoreceptors and its effect on visual detection. Journal of Neuroscience, 32(10): 3552–3562, 2012. ISSN 02706474. doi: 10.1523/JNEUROSCI.2144-11.2012.

49. Samuel M Wu and X. L. Yang. Electrical coupling between rods and cones in the tiger salamander retina. Proceedings of the National Academy of Sciences of the United States of America, 85(1): 275–278, 1988. ISSN 00278424. doi: 10.1073/pnas.85.1.275.

50. Nange Jin and Christophe P. Ribelayga. Direct evidence for daily plasticity of electrical coupling between rod photoreceptors in the mammalian retina. Journal of Neuroscience, 36(1):178–184, jan 2016. ISSN 15292401. doi: 10.1523/JNEUROSCI.3301-15.2016.

51. Jennifer J. O’Brien, Xiaoming Chen, Peter R Macleish, John O’Brien, and Stephen C Massey. Photoreceptor coupling mediated by connexin36 in the primate retina. Journal of Neuroscience, 32(13): 4675–4687, 2012. ISSN 02706474. doi: 10.1523/JNEUROSCI.4749-11.2012.

52. Sammy C.S. Lee, Paul R. Martin, and Ulrike Grünert. Topography of neurons in the rod pathway of human retina. Investigative Ophthalmology and Visual Science, 60(8):2848–2859, 2019. ISSN 15525783. doi: 10.1167/iovs.19-27217.

53. Yoshihiko Tsukamoto, K Morigiwa, M Ueda, and Peter Sterling. Microcircuits for night vision in mouse retina. The Journal of neuroscience : the official journal of the Society for Neuroscience, 21(21):8616–23, nov 2001. ISSN 1529-2401. doi: 10.1523/JNEUROSCI.21-21-08616.2001.

54. Denis A. Baylor, B. J. Nunn, and Julie L. Schnapf. The photocurrent, noise and spectral sensitivity of rods of the monkey Macaca fascicularis. The Journal of Physiology, 357(1): 575–607, dec 1984. ISSN 14697793. doi: 10.1113/jphysiol.1984.sp015518.

55. M. C. W. van Rossum and Robert G. Smith. Noise removal at the rod synapse. Visual Neuroscience, 15(5): 809–821, 1998.

56. Brenna Krieger, Mu Qiao, David L. Rousso, Joshua R. Sanes, and Markus Meister. Four alpha ganglion cell types in mouse retina: Function, structure, and molecular signatures. PLoS ONE, 12(7), jul 2017. ISSN 19326203. doi: 10.1371/journal.pone.0180091.

57. Lina Smeds, Daisuke Takeshita, Tuomas Turunen, Jussi Tiihonen, Johan Westö, Nataliia Martyniuk, Aarni Seppänen, and Petri Ala-Laurila. Paradoxical Rules of Spike Train Decoding Revealed at the Sensitivity Limit of Vision. Neuron, 104(3):576–587.e11, 2019. ISSN 10974199. doi: 10.1016/j.neuron.2019.08.005.

58. Chang Jin Jeon, Enrica Strettoi, and Richard H Masland. The major cell populations of the mouse retina. Journal of Neuroscience, 18(21): 8936–8946, 1998. ISSN 02706474. doi: 10.1523/JNEUROSCI.18-21-08936.1998.

59. Alapakkam P. Sampath and Fred Rieke. Selective Transmission of Single Photon Responses by Saturation at the Rod-to-Rod Bipolar Synapse. Neuron, 41(3): 431–443, 2004. ISSN 08966273. doi: 10.1016/S0896-6273(04)00005-4.

60. Dmitry S. Lebedev, Alexei L. Byzov, and Victor I. Govardovskii. Photoreceptor coupling and boundary detection. Vision Research, 38(20): 3161–3169, 1998. ISSN 00426989. doi: 10.1016/S0042-6989(98)00017-0.

61. David Attwell, Salvador Borges, Samuel M. Wu, and Martin Wilson. Signal clipping by the rod output synapse. Nature, 328(6130): 522–524, 1987. ISSN 00280836. doi: 10.1038/328522a0.

62. Feng Pan, David L. Paul, Stewart A. Bloomfield, and Béla Völgyi. Connexin36 is required for gap junctional coupling of most ganglion cell subtypes in the mouse retina. J Comp Neurol, 518(6): 911–927, 2010. doi: doi:10.1002/cne.22254.

63. Yoshihiko Tsukamoto and Naoko Omi. Functional allocation of synaptic contacts in microcircuits from rods via rod bipolar to AII amacrine cells in the mouse retina. Journal of Comparative Neurology, 521(15): 3541–3555, 2013. ISSN 00219967. doi: 10.1002/cne.23370.

64. Corinne Beier, Ulisse Bocchero, Lior Levy, Zhijing Zhang, Nange Jin, Stephen C. Massey, Christophe P. Ribelayga, Kirill Martemyanov, Samer Hattar, and Johan Pahlberg. Divergent outer retinal circuits drive image and non-image visual behaviors. Cell Reports, 39(13):111003, jun 2022. ISSN 22111247. doi: 10.1016/J.CELREP.2022.111003.

65. Markku Kilpeläinen, Johan Westö, Jussi Tiihonen, Anton Laihi, Daisuke Takeshita, Fred Rieke, and Petri Ala-Laurila. Primate retina trades single-photon detection for high-fidelity contrast encoding. Nature Communications, 15(1): 1–9, 2024. ISSN 20411723. doi: 10.1038/s41467-024-48750-y.

66. Alapakkam P. Sampath, Katherine J. Strissel, Rajesh Elias, Vadim Y. Arshavsky, James F. McGinnis, Jeannie Chen, Satoru Kawamura, Fred Rieke, and James B. Hurley. Recoverin improves rod-mediated vision by enhancing signal transmission in the mouse retina. Neuron, 46(3): 413–420, 2005. ISSN 08966273. doi: 10.1016/j.neuron.2005.04.006.

67. Rebecca Holmes, Michelle Victora, Ranxiao Frances Wang, and Paul G. Kwiat. Measuring temporal summation in visual detection with a single-photon source. Vision Research, 140: 33–43, 2017. ISSN 18785646. doi: 10.1016/j.visres.2017.06.011.

68. William W. Dawson and Joseph M. Harrison. Bloch’s law for brief flashes of large angular subtense. Perceptual and Motor Skills, 36: 1055–1061, 1973.

69. Yu Zheng Le, John D. Ash, Muayyad R. Al-Ubaidi, Ying Chen, Jian Xing Ma, and Robert E. Anderson. Targeted expression of Cre recombinase to cone photoreceptors in transgenic mice. Molecular Vision, 10: 1011–1018, 2004. ISSN 10900535.

70. Patrick H. Roseboom, M. a.Aryan Namboodiri, Drazen B. Zimonjic, Nicholas C. Popescu, Ignacio R. Rodriguez, Jonathan A. Gastel, and David C. Klein. Natural melatonin ‘knockdown’ in C57BL/6J mice: Rare mechanism truncates serotonin N-acetyltransferase. Molecular Brain Research, 63(1): 189–197, 1998. ISSN 0169328X. doi: 10.1016/S0169-328X(98)00273-3.

71. Yumiko Umino, Rose Pasquale, and Eduardo Solessio. Visual temporal contrast sensitivity in the behaving mouse shares fundamental properties with human psychophysics. eNeuro, 5(4): 1–14, 2018. ISSN 23732822. doi: 10.1523/ENEURO.0181-18.2018.

72. Neil A. Macmillan and C. Douglas Creelman. Detection Theory: A User’s Guide. Lawrence Erlbaum Associates, New York, 2nd edition, 2005. ISBN 0805842306.

73. John A. Swets, Wilson P. Tanner, and Theodore G. Birdsall. Decision Processes In Perception. Psychological Review, 68(5): 301–340, 1961. ISSN 0033295X. doi: 10.1037/h0040547.

74. Michael J. Hautus. Corrections for extreme proportions and their biasing effects on estimated values of d’. Behavior Research Methods, Instruments, & Computers, 27(1): 46–51, 1995. ISSN 07433808. doi: 10.3758/BF03203619.

75. Arkady L. Lyubarsky, Lauren L. Daniele, and Edward N. Pugh. From candelas to photoisomerizations in the mouse eye by rhodopsin bleaching in situ and the light-rearing dependence of the major components of the mouse ERG. Vision Research, 44(28 SPEC.ISS.):3235–3251, 2004. ISSN 00426989. doi: 10.1016/j.visres.2004.09.019.

76. Mark Bushnell, Yumiko Umino, and Eduardo Solessio. A system to measure the pupil response to steady lights in freely behaving mice. Journal of Neuroscience Methods, 273(315): 74–85, 2017. doi: 10.1016/j.jneumeth.2016.08.001.A.

77. Horace B. Barlow, W R Levick, and M Yoon. Responses to Single Quanta of Light in Retinal Ganglion Cells of the Cat. Vision Research, 11(3): 87–101, 1971.

78. Denis A. Baylor, G. Matthews, and King Wai Yau. Two components of electrical dark noise in toad retinal rod outer segments. The Journal of Physiology, 309(1):591–621, dec 1980. ISSN 14697793. doi: 10.1113/jphysiol.1980.sp013529.

79. Horace B. Barlow. Retinal and central factors in human vision limited by noise. In Horace B. Barlow and P. Fatts, editors, Photoreception in Vertebrates, chapter 19, pages 337–358. Academic Press, London, 1977.

80. Greg D. Field, Alapakkam P. Sampath, and Fred Rieke. Retinal Processing Near Absolute Threshold: From Behavior to Mechanism. Annu. Rev. Physiol, 67: 491–514, 2005. doi: 10.1146/annurev.physiol.67.031103.151256.

81. Theodore E. Cohn, Daniel G. Green, and Wilson P. Tanner. Receiver operating characteristic analysis : Application to the study of quantum fluctuation effects in optic nerve of rana pipiens. Journal of General Physiology, 66(5):583–616, 1975. ISSN 15407748. doi: 10.1085/jgp.66.5.583.

82. Larry N. Thibos, W. R. Levick, and Theodore E. Cohn. Receiver Operating Characteristic Curves for Poisson Signals. Biological Cybernetics, 33: 57–61, 1979.

83. Christian Kaernbach. Theory and Evaluative Reviews Poisson signal-detection theory : Link between threshold models. Perception & Psychophysics, 50(5): 498–506, 1991.

84. Frederick A.A. Kingdom and Nicolaas Prins. Psychophysics: A Practical Introduction. Academic Press, London, 1st edition, 2010.

85. Haruhisa Okawa and Alapakkam P. Sampath. Temporal Transformation of the Rod Single-Photon Response in the Retinal Circuitry. Investigative Ophthalmology and Visual Science, 66(2), 2025. ISSN 15525783. doi: 10.1167/iovs.66.2.52.

86. Andrew B. Watson. Probability Summation Over Time. Vision Research, 19: 515–522, 1979.

87. Kenneth P. Burnham and David R. Anderson. Model Selection and Inference: A Practical Information-Theoretic Approach. Springer-Verlag, New York, 2nd edition, 2002. ISBN 0387953647. doi: 10.2307/3803117.

88. M. Christopher Newland. An Information Theoretic Approach to Model Selection: A Tutorial with Monte Carlo Confirmation. Perspectives on Behavior Science, 42 (3):583–616, 2019. ISSN 25208977. doi: 10.1007/s40614-019-00206-1.

89. Galit Shmueli. To explain or to predict? Statistical Science, 25(3): 289–310, 2010. ISSN 08834237. doi: 10.1214/10-STS330.

90. Elliott Sober. Instrumentalism, Parsimony, and the Akaike Framework. Philosophy of Science, 69(S3):S112–S123, 2002. ISSN 0031-8248. doi: 10.1086/341839.

91. Eric Jan Wagenmakers and Simon Farrell. AIC model selection using Akaike weights. Psychonomic Bulletin and Review, 11(1): 192–196, 2004. ISSN 10699384. doi: 10.3758/BF03206482.

92. Bradley Efron and Robert J. Tibshirani. An Introduction to the Bootstrap. Chapman & Hall/CRC, Boca Raton, 1994.

93. SAS Institute Inc. SAS/STAT® 15.3 User’s Guide. SAS Institute Inc., Cary, NC, 2023.

94. Arielle S. Selya, Jennifer S. Rose, Lisa C. Dierker, Donald Hedeker, and Robin J. Mermelstein. A practical guide to calculating Cohen’s f2, a measure of local effect size, from PROC MIXED. Frontiers in Psychology, 3(APR):1–6, 2012. ISSN 16641078. doi: 10.3389/fpsyg.2012.00111.

95. Jacob Cohen. Statistical Power Analysis for the Behavioral Sciences Second Edition. Lawrence Erlbaum Associates, 1988. ISBN 0805802835.

