## Supplemental Information for "Retinal Cx36 gap junctions in the inner and outer plexiform layers differentially control visual thresholds"

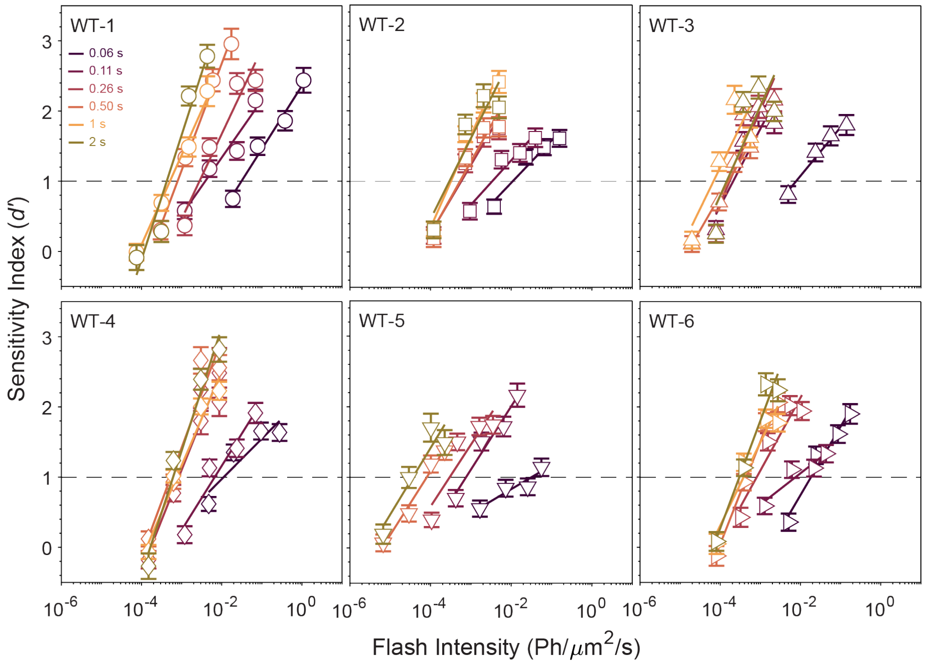


Figure S1.1: $d'$ values with best fit psychometric functions (Eq. 3) for all WT mice, at each duration tested.

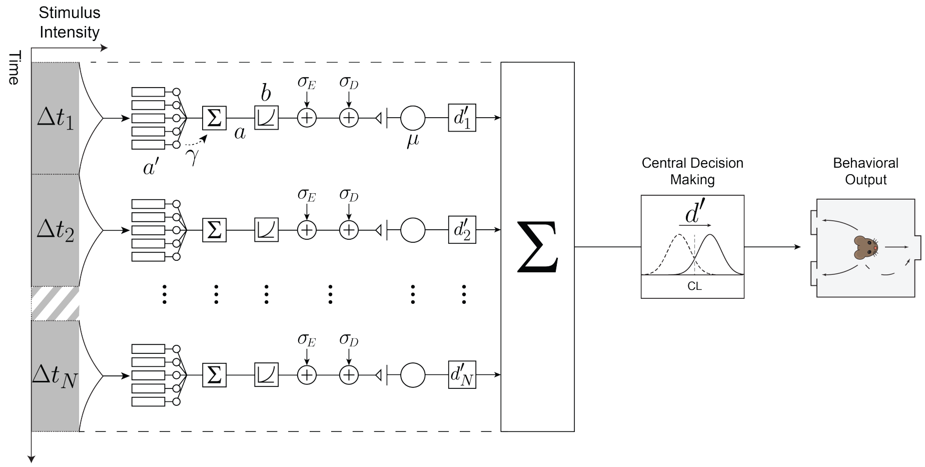


Figure S1.2: **Diagrammatic outline of the Signal Detection Theoretic model of temporal summation**. We model the retina as consisting of a series of parallel detectors, that integrate photons spatially over an array of rods (the number of which is represented by the $a'$ parameter) and within an integration time ($\Delta t_i$). The resultant signal undergoes a compressive non-linearity ($b$) which gives rise to a signal $\mu$ at the level of the detector, which we propose may be a ganglion cell. Note, only a fraction of the responses generated by the rods will are integrated (given by $\gamma$). Additionally, there is both external ($\sigma_E$) and internal ($\sigma_D$) noise that is added to the signal. This results in a signal-to-noise ratio $d'_i$ at the level of the detector output. As the stimulus is prolonged, the number ($N$) of detectors responding to the stimulus increases linearly. The output from each of these N detectors are then subject to a second order summation. This results in a "total" $d'$ at the level of central decision making, which, along with a certain criterion (CL), the observer uses to determine their response.

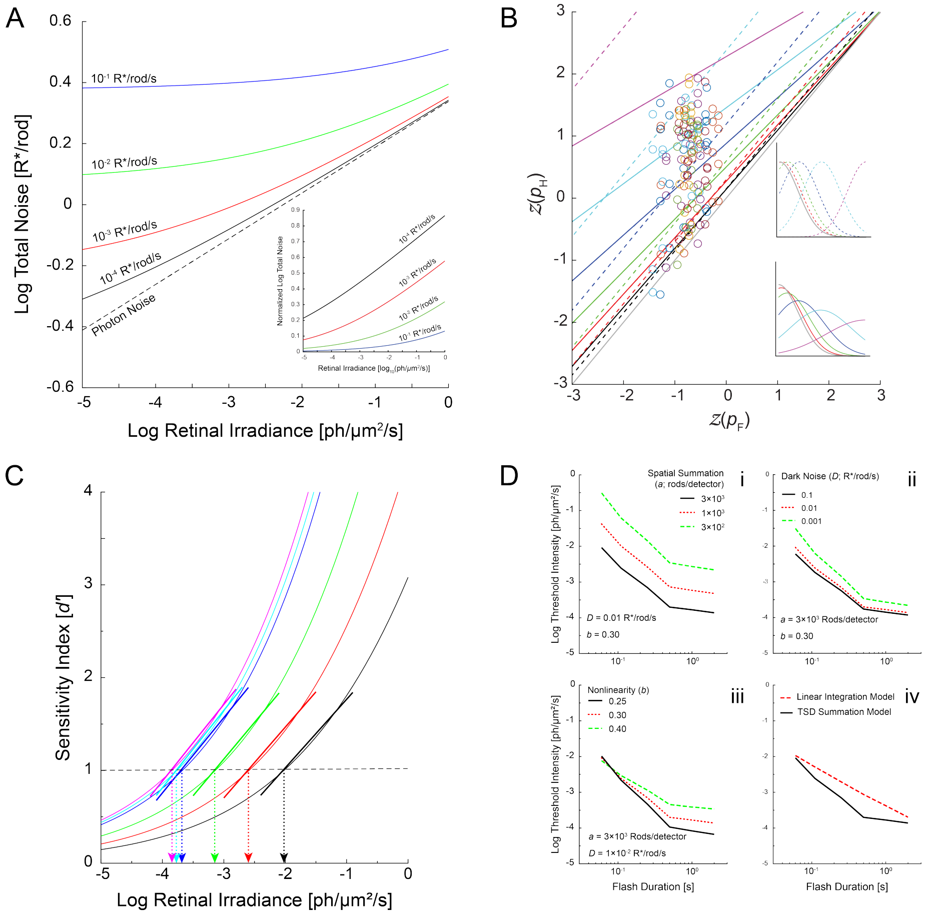


Figure S1.3: **Properties of the temporal summation model.** **A** Total noise plotted as a function of retinal irradiance with no internal dark noise, and with increasing amounts of dark noise (in equivalent R*/rod/s). Inset shows total noise normalized to its minimum as a function of retinal irradiance. **B** Receiver operating characteristic curves (ROC) on normal deviate axes, where $z(p_H)$ is plotted as a function of $z(p_F)$ for a range of stimulus intensities. $z$ROC curves assuming equal variance (see top inset) are shown as dashed lines. $z$ROCs assuming variance increases with stimulus intensity (see bottom inset) are shown as solid lines, where it is assumed $a = 3{,}000$ and $b = 0.30$. Grey solid line represents baseline internal noise. Also plotted are the $z(p_H)$, $z(p_F)$ pairs for each WT mouse. **C** Sensitivity indices predicted by the TSD model for $N = 1$, 2, 4, 8, 8, and 8 (thin solid lines) as well as log-linear psychometric functions (thick solid lines) fit to the simulated data using Equation 3. **D** Effects of changing $a$ (i), $D$ (ii), and $b$ (iii) on model-predicted t.v.d. curves, as well as comparison of the TSD summation model with the linear integration model (iv).

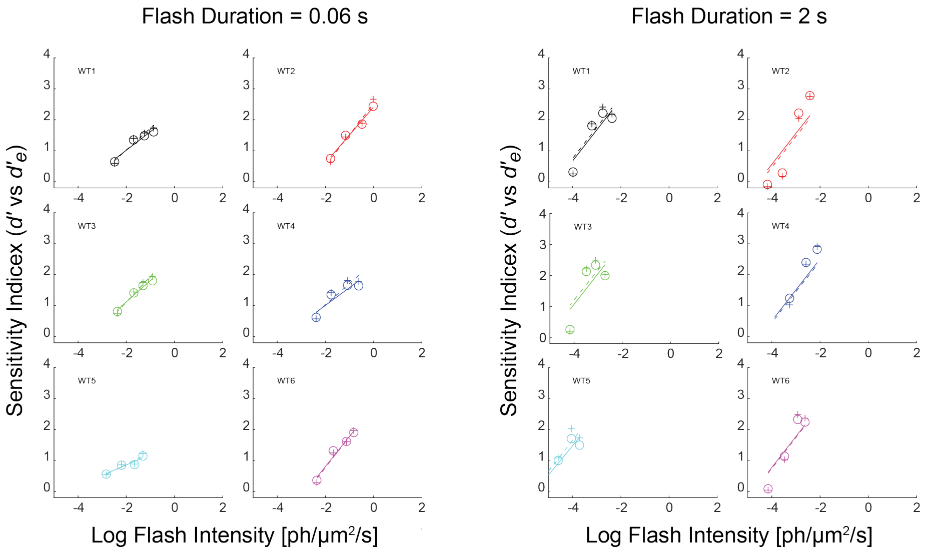


Figure S1.4: $d'$ **and** $d'_e$ **lead to similar threshold values.** $d'$ (circles) and $d'_e$ (crosses) values as a function of log flash intensity for all 6 WT mice at $\Delta t = 0.06$ s and $\Delta t = 2$ s. The $d'_e$ values are calculated assuming $a = 2{,}886$ and $b = 0.30$. Additionally, best-fit psychometric functions (solid lines are $d'$ ; dotted lines are $d'_e$) are plotted for $d'$ and $d'_e$ values.

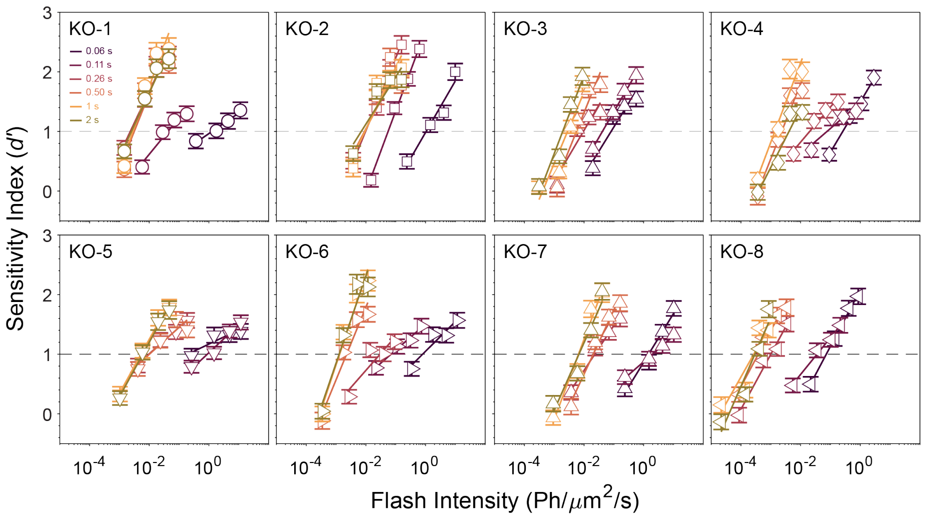


Figure S2.1: $d'$ values with best fit psychometric functions (Eq. 3) for all Cx36KO mice, at each duration tested.

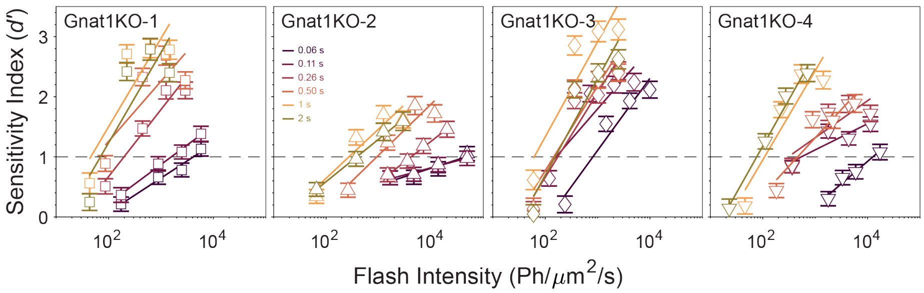


Figure S2.2: $d'$ values with best fit psychometric functions (Eq. 3) for all Gnat1KO mice, at each duration tested.

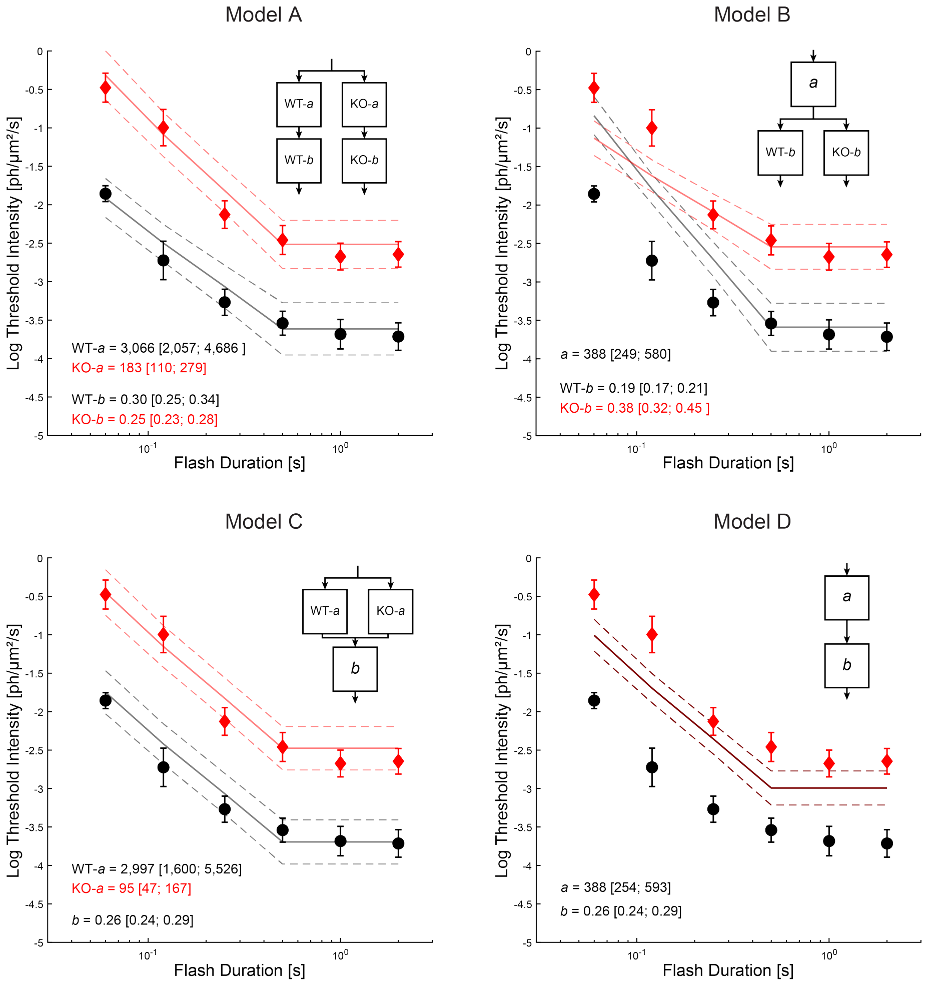


Figure S2.3: Average t.v.d. data for WT (black circles) and Cx36KO (red diamonds) mice and each of the four candidate models with best-fit parameters. Block diagrams illustrating each model are given in insets.

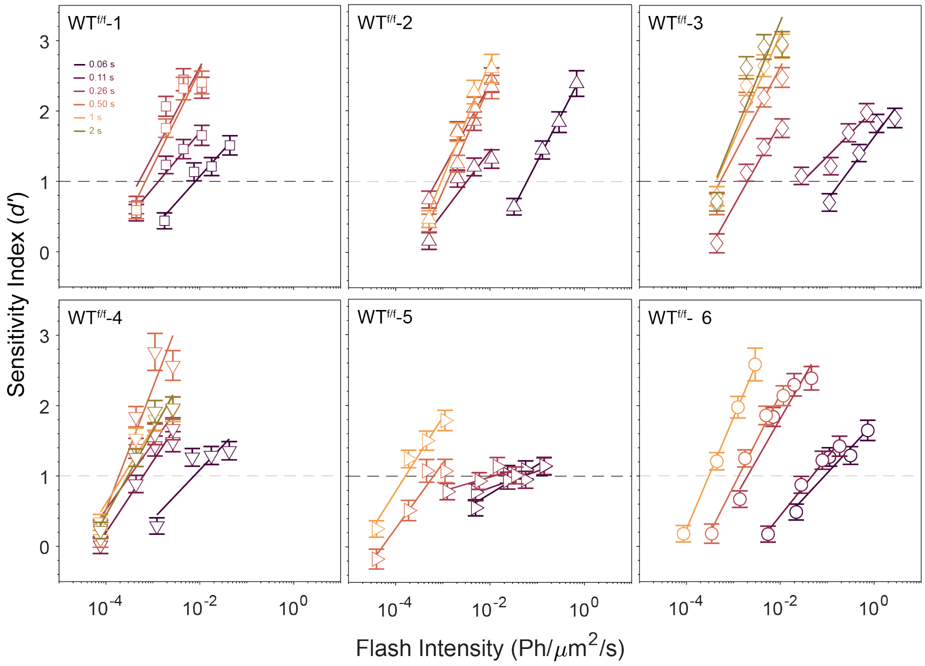


Figure S3.1: $d'$ values with best fit psychometric functions (Eq. 3) for all WT$^{f/f}$mice, at each duration tested.

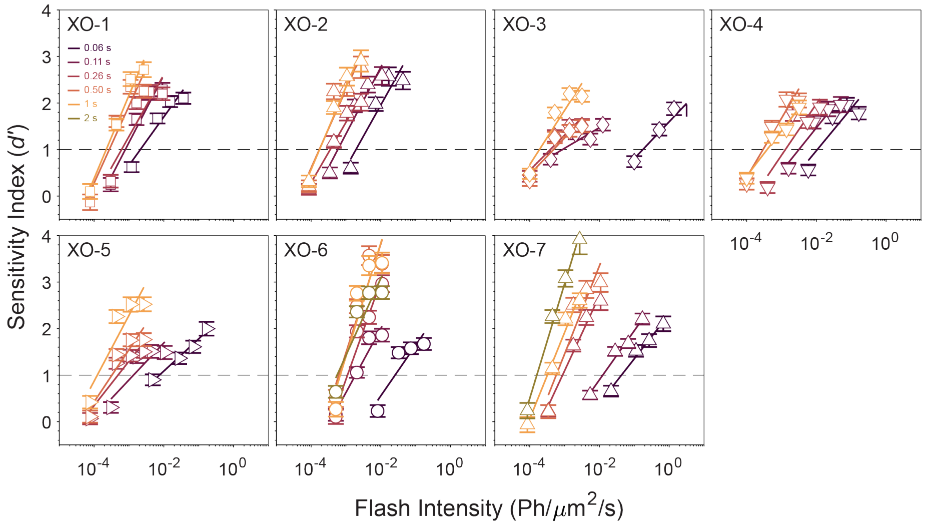


Figure S3.2: $d'$ values with best fit psychometric functions (Eq. 3) for all Cx36XO mice, at each duration tested.

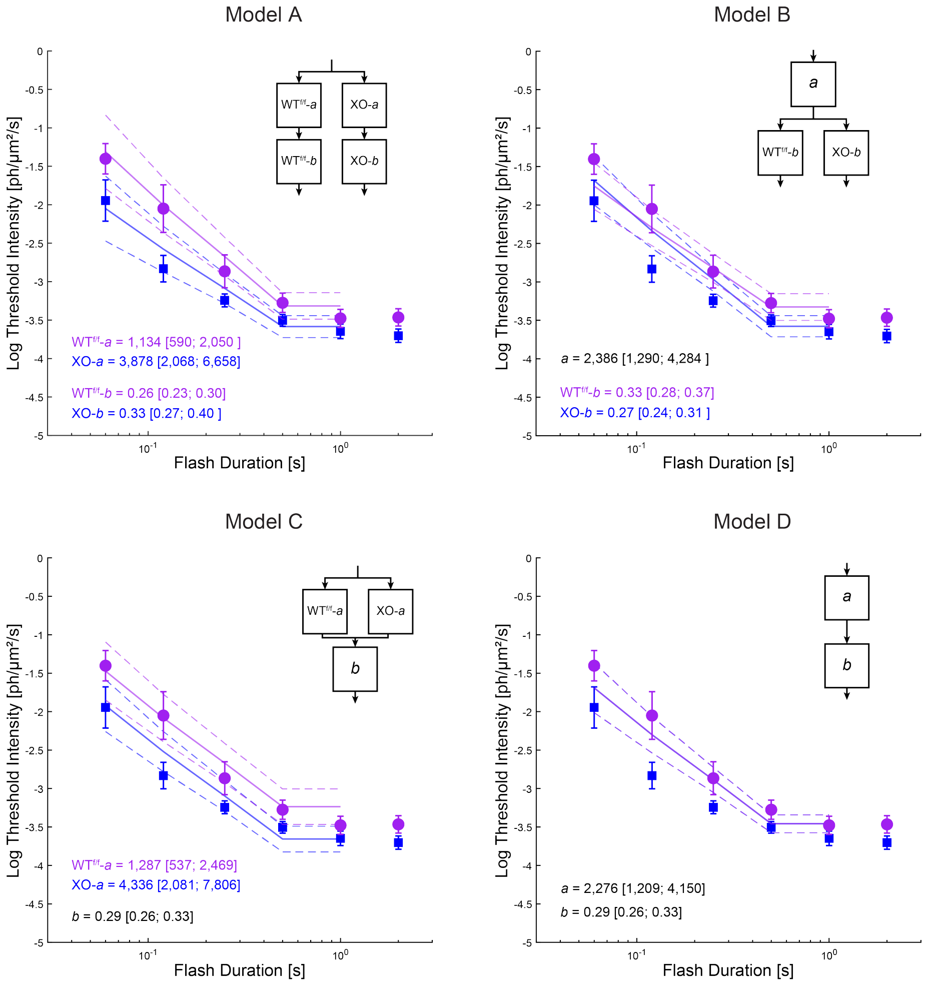


Figure S3.3: Average t.v.d. data for WT$^{f/f}$(purple circles) and Cx36XO (blue squares) mice and each of the four candidate models with best-fit parameters. Block diagrams illustrating each model are given in insets.
